# Transient disruption of the inferior parietal lobule impairs the ability to attribute intention to action

**DOI:** 10.1101/2020.04.18.047787

**Authors:** Jean-François Patri, Andrea Cavallo, Kiri Pullar, Marco Soriano, Martina Valente, Atesh Koul, Alessio Avenanti, Stefano Panzeri, Cristina Becchio

## Abstract

Although it is well established that fronto-parietal regions are active during action observation, whether they play a causal role in the ability to infer others’ intentions from visual kinematics remains undetermined. In experiments reported here, we combined offline continuous theta-burst stimulation (cTBS) with computational modeling to reveal single-trial computations in the inferior parietal lobule (IPL) and inferior frontal gyrus (IFG). Participants received cTBS over the left anterior IPL and the left IFG pars orbitalis, in separate sessions, before completing an intention discrimination task (discriminate intention of observed reach-to-grasp acts) or a kinematic discrimination task (discriminate peak wrist height of the same acts) unrelated to intention. We targeted intentions-sensitive regions whose fMRI-measured activity accurately discriminated intention from the same action stimuli. We found that transient disruption of activity of the left IPL, but not the IFG, impaired the observer’s ability to judge intention from movement kinematics. Kinematic discrimination unrelated to intention, in contrast, was largely unaffected. Computational analyses revealed that IPL cTBS did not impair the ability to ‘see’ changes in movement kinematics, nor did it alter the weight given to informative versus non-informative kinematic features. Rather, it selectively impaired the ability to link variations in informative features to the correct intention. These results provide the first causal evidence that left anterior IPL maps kinematics to intentions.

## Introduction

When watching others in action, we readily infer their intentions from subtle changes in the way they move. Theoretical work [1–5] and related experimental findings (e.g., [6–9]) suggest that the ability to read the intention of an observed action is mediated by the fronto-parietal action observation network. Despite two decades of research, however, the specific neural computations involved in this ability remain unclear and causally untested [10].

A major difficulty in studying how intentions are inferred from others’ actions is the ever-changing nature of movement kinematics [11,12]. Movement is “repetition without repetition” [13]. Averaging across repeats of nominally identical, but actually different motor acts, as done in standard trial-averaged analyses, can obscure how intention information is encoded in trial-to-trial variations in movement kinematics [14]. More importantly, the brain does not operate according to an average response over averaged kinematics. Real-world intention attribution requires real-time readout of intention-information encoded within a specific motor act. Thus, studying intention attribution with single-trial resolution is critical for understanding how intention readout maps to the multiplicity and variability of kinematic patterns.

Here, we developed a novel analysis framework to uncover information from single-trial level kinematics. This framework was inspired by recent mathematical advances in determining how sensory information encoded in a neural population is read out to inform single-trial behavioral choice [15–17]. In this study, we adapted this approach to investigate neural computations performed in the left inferior parietal lobule (IPL), a core region of the action observation network, and explore the hypothesis that neural computations of the left IPL play a causal role in the attribution of intention to action.

Activity in IPL is related to the coding of intention in both monkeys and humans [3,6,7,9,18,19]. In monkeys, the PFG area, found on the convex of the IPL, contains visuo-motor neurons whose activity is modulated by the inferred intention of an observed act (e.g., grasp-to-eat) [6,7]. In humans, intentions inferred from a set of observed actions can be decoded using spatial patterns of activity in the left IPL [9]. These results suggest that the left IPL contains information about intentions of observed actions. However, the level of causal inference afforded by observation of neural activity is limited [20,21]. This is because, in the absence of perturbation, the relationship between neural information and behavior remains correlational. Moreover, simple observation of neural activity cannot determine what features of neural representations are read out by other regions and what neural computations affect downstream processing [15,22,23]. The contribution of IPL to intention attribution, both in terms of function – *does IPL plays a causal role in intention to action attribution?* – and content – *what and how does IPL compute?* – remains therefore largely undefined [24].

To investigate these questions, we applied continuous theta-burst transcranial magnetic stimulation (cTBS) to reversibly reduce cortical excitability in two fronto-parietal sites within the action observation network [25–29]. Specifically, we targeted two intentions-sensitive regions within the left IPL and the left inferior frontal gyrus (IFG) determined based on multi-voxel pattern analysis of fMRI data [9].

We investigated how transient disruption of activity in these regions influences the observer’s readout computations involved in extracting intention-related information from movement kinematics. Single-trial analyses combined with a set of task manipulations revealed that disruption of activity in the IPL site, but not the IFG site, impaired an observer’s ability to interpret the intentional significance of discriminative kinematic features.

## Results

### Causal contribution of IPL to intention discrimination

To perturb fronto-parietal sites within the action observation network, we used a cTBS protocol delivered offline for 40 s [30–32]. In three separate sessions, participants either received no cTBS or Magnetic Resonance Imaging (MRI)-guided cTBS to the left IPL or left IFG before completing a two-alternative, forced-choice (2AFC) discrimination of intention (Fig. 1a-c).

**Fig. 1.**
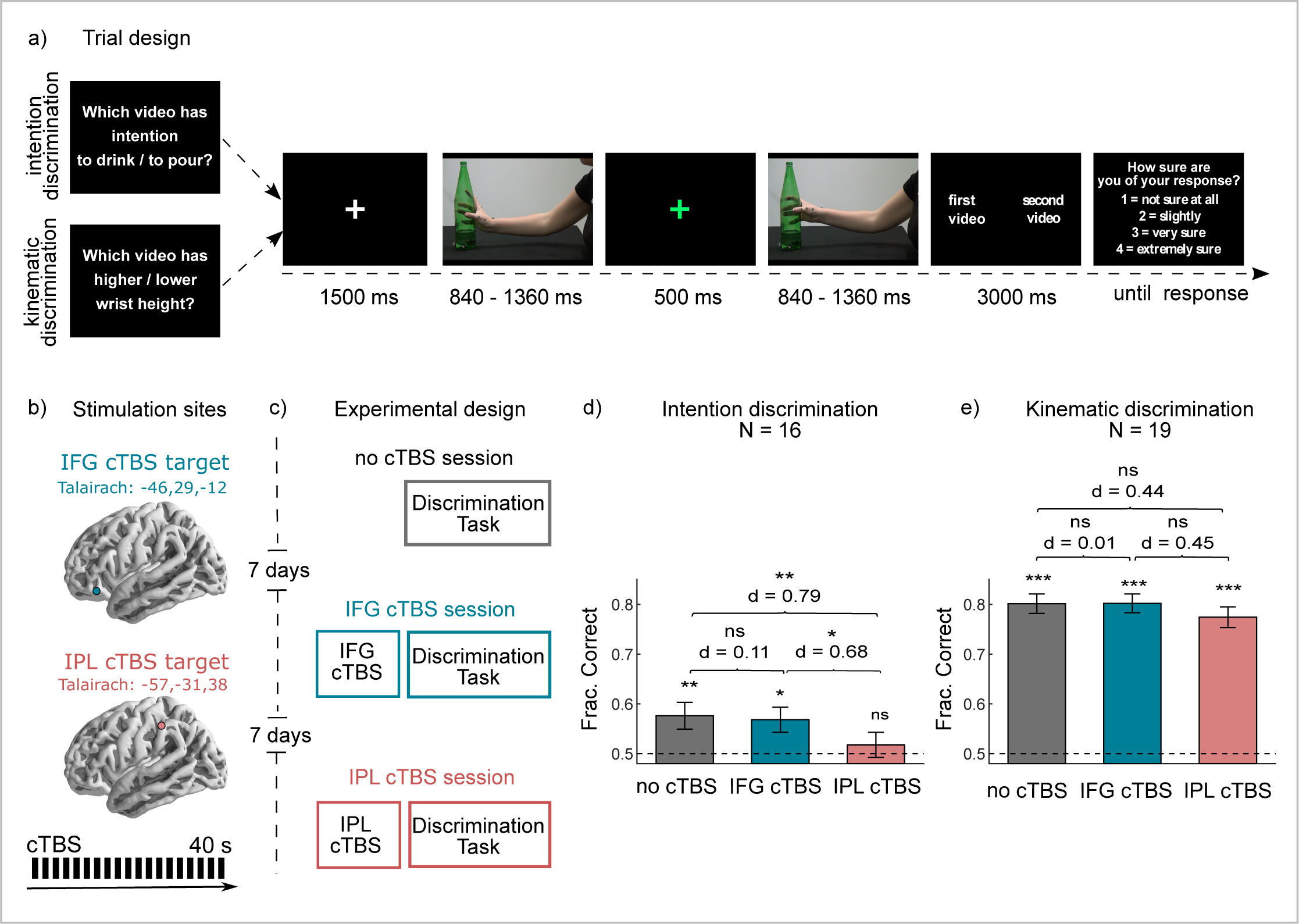
Experimental design and behavioral discrimination results. **a** Trial design of the discrimination tasks. **b** cTBS targets and MRI-guided cTBS protocol. **c** Sketch of the experimental design. **d-e** Discrimination performance (fraction correct) in the intention discrimination task and in the kinematic discrimination task. Histograms represent mean ± SEM across participants. Number of participants in each task (N) and Cohen’s effect size (d) of each comparison are reported.

To capture natural movement variability, we selected 120 representative reach-to-grasp acts – 60 for each intention – from a large dataset obtained by filming naïve participants reaching toward and grasping a bottle with the intent to drink or to pour. Each trial displayed two reach-to-grasp acts in two consecutive temporal intervals: one interval contained a grasp-to-pour act, the other interval a grasp-to-drink act. Participants (N = 16) were required to indicate, on each trial, the interval displaying the reach-to-grasp performed with the intent to drink (or to pour; see Methods), and, at the end of the trial, to rate the confidence of their choice (Fig. 1a).

Individual targets for cTBS were determined based on a re-analysis of published fMRI data [9] recorded during an action observation task using the same action stimuli employed in the current study. We targeted intentions-sensitive regions in the left anterior IPL and the left IFG pars orbitalis whose activity could accurately discriminate one intention from the other (see Methods and Supplementary Fig. 1a).

We used Logistic Mixed Effects Models [33] (LMEM, see Methods) to test statistically whether average discrimination performance differed from chance and across sessions. Discrimination performance was computed as the fraction of correct responses, given that there was no response bias (Supplementary Fig. 1b). Discrimination performance was higher than chance in no cTBS and IFG cTBS, but did not differ from chance post IPL cTBS (Fig. 1d). IPL cTBS resulted in significantly reduced intention discrimination relative to both no cTBS and IFG cTBS, with effect size Cohen’s d = 0.79 and 0.68, respectively (Fig. 1d). The effect of IFG cTBS relative to no cTBS, in contrast, was not significant (Fig. 1d). This suggests that cTBS to IPL, but not to IFG, impaired ability to discriminate intention.

### Discrimination of individual kinematic features following cTBS

To investigate the selectivity of the above reported effects to intention discrimination, we tested a new cohort of participants (N = 19) carrying out a 2AFC kinematic discrimination task unrelated to intention. Action stimuli and task parameters were identical to that of the intention discrimination task except that participants were required to discriminate differences in the peak wrist height of the observed acts. As shown in Fig. 1e, despite the larger number of participants and thus greater statistical power when compared to the intention task, the effect of cTBS to IPL (or to IFG) on kinematic discrimination performance was smaller and not significant (Fig. 1e). This suggests that following IPL and IFG cTBS observers retained to a large extent the ability to detect changes in individual kinematic features when instructed to do so.

We were concerned that lack of cTBS effects on kinematic discrimination might be related to the relative ease of the kinematic task. To control for task difficulty, we ranked trials in each task based on intention-related information encoded in movement kinematics (see Supplementary Fig. 1c) and restricted the analyses to the 10% most informative trials (n = 9) for intention discrimination and the 10% least informative trials (n = 9) for kinematic discrimination. With this selection of trials, kinematic discrimination performance did not differ from intention discrimination performance under no cTBS. Even in this control case, we found that, consistent with results utilizing all trials, the effect of IPL cTBS relative to both no cTBS and IFG cTBS on intention discrimination persisted with large effect size. In contrast, cTBS (to either IPL or IFG) had no significant effect on kinematic discrimination even with performance-matched trial subsampling. Collectively, these analyses suggest that IPL cTBS did not impair the ability to ‘see’ changes in individual kinematic features.

### Using logistic regression to relate intention encoding and readout at the single-trial level

Having demonstrated that post IPL cTBS observers retain the ability to process individual kinematic features, we next employed logistic regression to quantify intention encoding – the mapping, on each trial, of intention to movement kinematics – and intention readout – the mapping of movement kinematics to intention choice.

We hypothesized that discrimination performance can be explained as the intersection, at the single trial level, between intention encoding and intention readout [15,34]. We also hypothesized that IPL cTBS impacts on correct intention readout, that is, how well intention information encoded in movement kinematics is read out to inform intention choice.

### Encoding of intention-related information

To obtain a measure of intention-related information encoded in movement kinematics (of which example single trial traces are shown in Fig. 2a, b), we quantified single-trial variations in movement kinematics as a 64-dimensional kinematic vector reporting the values of all kinematic variables over time (16 kinematic variables at 4 time epochs) and used it as predictor in a logistic regression of the intention of the observed reach-to-grasp acts. For each trial, the encoding model computes the probability of the first interval to display a grasp-to-drink act (and thus of the second interval to display a grasp-to-pour act) as a combination of the features of the kinematic vector for that trial [35] (Fig. 2c, d). Intention encoding can thus be described by the set of regression coefficients across kinematic features.

**Fig. 2.**
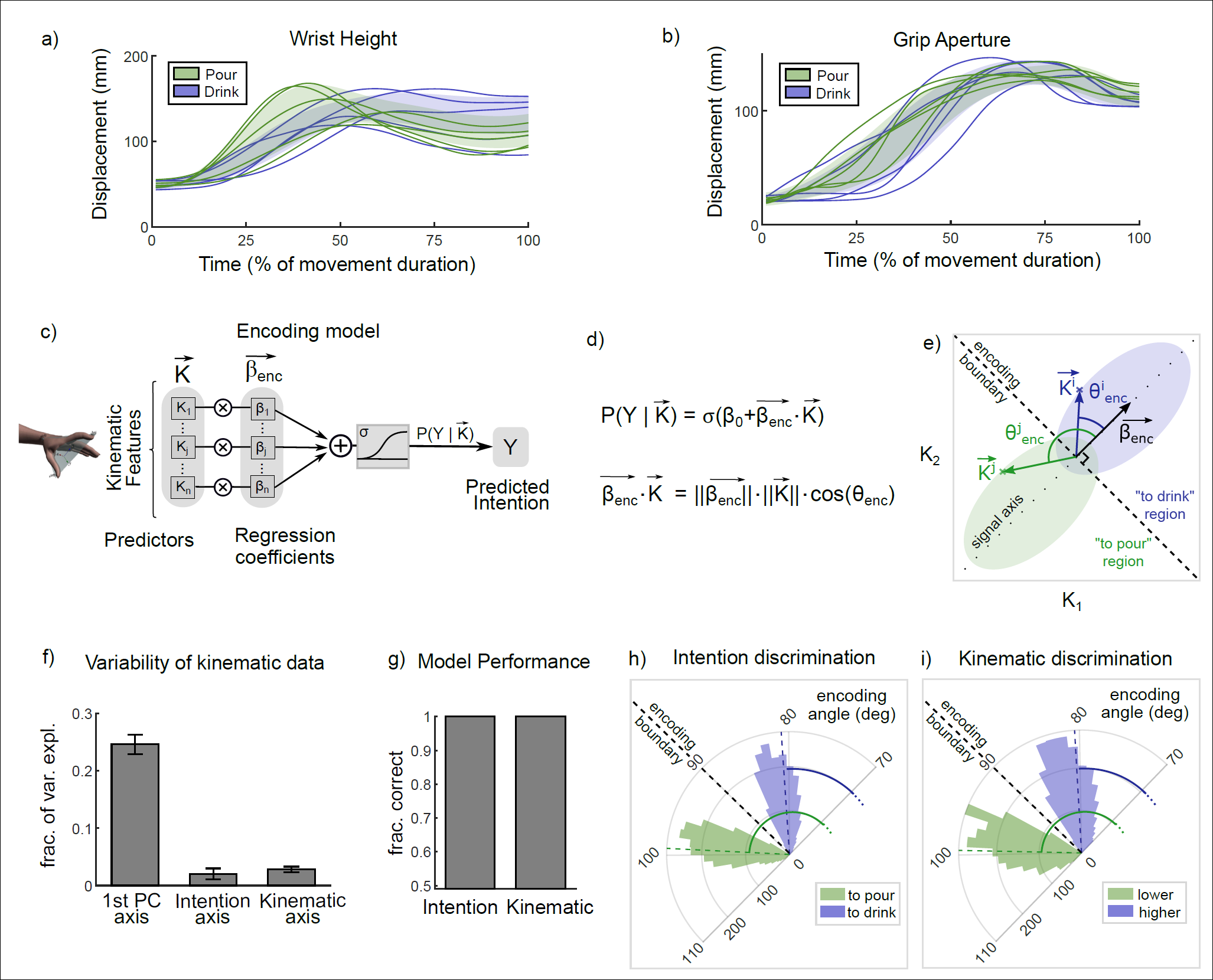
Encoding of task discriminant information in movement kinematics. **a-b** Time course of wrist height and grip aperture for reach-to-grasp acts performed with the intent to drink and to pour. Colored curves display representative trajectories for each intention, colored areas display one standard deviation across executed trials. **c-d** Schematic of the encoding model. Block diagram representation (c) and equation (d) of the logistic regression used to quantify intention information in movement kinematics. **e** Sketch of the encoding model in a simplified kinematic space spanning only two kinematic features K_1_, K_2_). The two elliptic regions represent the intention conditional probability distributions of the two features. The encoding boundary optimally separates the kinematics patterns into ‘to drink’ and ‘to pour’ regions. The encoding vector 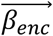 indicates the maximally discriminative axis. The angle *θ*_*enc*_ between the encoding vector and the single-trial kinematic vector 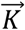 can be used to classify single trials according to intention. Two different single-trial kinematic vectors with superscript *i* and *j* are shown. **f** Fraction of variance explained across executed acts computed along three different kinematic axes (1^st^ PC axis of largest variance, the axis of largest intention discrimination, and the axis of largest wrist height discrimination). **g** Performance of encoding models, quantified as the fraction of correctly predicted trials. **h-i** Distribution of encoding angles across trials in the intention discrimination task (h) and in the kinematic discrimination task (i). For graphical representation, the 70-110° angle range is expanded to a semi-circle. In panels (f)-(g), histograms represent mean ± SEM.

Fig. 2e shows a sketch of the encoding model in a hypothetical, simplified kinematic space spanning only two kinematic features. The encoding boundary defines the border that best separates the kinematic patterns of the two intentions. The encoding vector, orthogonal to the encoding boundary, indicates the information axis along which changes in kinematics maximally discriminate between intentions. Single-trial kinematic vectors are classified as ‘to pour’ or ‘to drink’ depending on which side of the boundary they fell or, equivalently, according to the angle they formed with the encoding vector. Since in our convention the encoding vector points towards ‘to drink,’ 0-90° encoding angles indicate ‘to drink,’ whereas 90-180° encoding angles indicate ‘to pour.’

Kinematic features were variable across individual trials, with only a small amount of variance (about 3%) aligned along the encoding vector and thus available for intention discrimination (Fig. 2f). Despite the small amount of intention-related signal hidden within the highly variable kinematic data, our encoding model was able to decode intention with perfect accuracy (Fig. 2g, h). This indicates that intention-related variation in grasping kinematics, although small, is nevertheless sufficient to fully specify intention-information in each trial. As expected by task design, we also observed a perfect decoding accuracy for the kinematic discrimination task (Fig. 2g, i).

### Readout of intention-related information

Having quantified intention information encoded in single-trial kinematics, we next asked how human observers read out such information. To assess this, we used logistic regression to predict the probability of intention choice in each trial as a combination of the features of the kinematic vector for that trial (Fig. 3a-c). Regression coefficients were estimated separately for each participant for no cTBS, IPL cTBS, and IFG cTBS sessions.

**Fig. 3.**
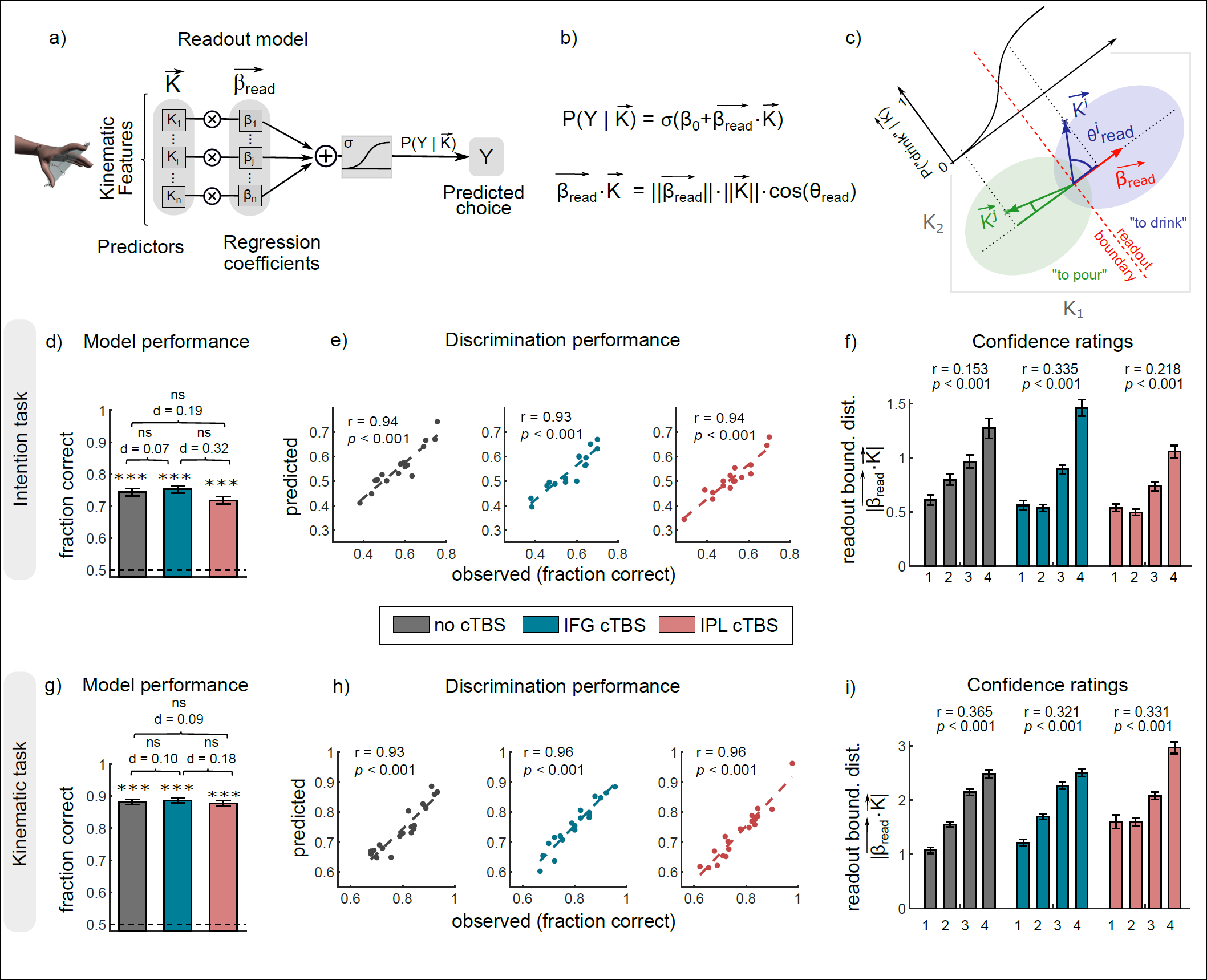
Readout model of intention discrimination from single-trial kinematics. **a-b** Readout model. Block diagram representation (a) and equation (b) of the logistic regression used to quantify intention information readout. **c** Sketch of the readout model in a simplified kinematic space spanning only two kinematic features. The two elliptic regions represent the probability distributions of the two features conditional to intention choice. The readout vector 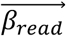 of regression coefficients indicates the direction in feature space that maximally discriminates observers’ choices. The direction and distance of the single-trial kinematic vector from the readout boundary determines, through the sigmoid logistic function, the probability of intention choice in that trial. **d** Performance of the readout model in predicting observers’ choices in the intention discrimination task quantified as fraction correct. **e** Scatterplot of the relationship between the observed discrimination performance and the one predicted by the readout model across individual participants in the intention discrimination task. **f** Distance of the single-trial kinematic vector from the readout boundary as a function of confidence ratings for the intention discrimination task. This distance was computed as modulus of the scalar product between single-trial kinematic vector and the readout vector. As shown in the panel (c), the larger this distance, the further from chance is the probability of intention choice predicted by the model. The green single-trial kinematic vector in panel (c), for example, has a larger distance and thus a larger probability of intention choice than the blue kinematic vector. **g-i** Same as (d)-(f) for the kinematic discrimination task. In panels (d),(g), Cohen’s effect size (d) of each comparison are reported.

We fitted the readout model, separately for each participant, to single-trial intention choices and used confidence ratings reported by observers for independent validation of the model. Across trials and participants, model performance, measured as the fraction of intention choices correctly predicted by the model, was significantly above chance (Fig. 3d).

Although confidence ratings were not used for fitting model parameters, we also found a positive, trial-to-trial relationship between the observer’s confidence in their intention choice and the distance of single-trial kinematics vector from the readout boundary – the border that best separates kinematic patterns for the two intention choices (Fig. 3f). This suggests that intention choices on trials farther away from the readout boundary (and thus classified with greater confidence by the model) were also endorsed with higher confidence by human observers. Similar results were obtained for a readout model using single-trial differences in movement kinematics to predict kinematic discrimination (Fig. 3g-i).

Collectively, the above analyses suggest that our readout model was able to capture task performance, providing a plausible description of how well and how confidently observers performed the discrimination tasks based on single-trial kinematics.

### Transient disruption of IPL does not decrease sensitivity of intention readout to movement kinematics

Having verified that our readout model could account for discrimination performance, we next used it to test alternative hypotheses regarding the functional effects of transient disruption of IPL on intention attribution. First, we considered the possibility that cTBS to IPL decreases the sensitivity of intention readout to single-trial variations in movement kinematics. If so, one would expect statistical dependency between single-trial kinematics and intention choices to be weaker following IPL cTBS. To test this formally, we compared the fraction of intention choices correctly predicted by the model across sessions. We found no difference between no cTBS, IPL cTBS and IFG cTBS; single-trial kinematics was equally predictive of intention choice across sessions (*p* = 0.60) (Fig. 3d). These analyses suggest that diminished sensitivity of intention readout to kinematics cannot account for the inferior discrimination performance following IPL cTBS.

### Transient disruption of IPL causes misalignment of intention readout with respect to encoding

Next, we asked whether IPL cTBS alters the ability to read out correctly intention-related information encoded in movement kinematics. An intuitive visualization of how well readout captures intention-related information in movement kinematics is provided by the angle between the encoding vector and the readout vector orthogonal to the readout boundary (Fig. 4a, b). The smaller the angle between these vectors, the larger the across-trial alignment between intention encoding and readout in kinematic space, and thus the larger the probability that intention-information is read out correctly across trials.

**Fig. 4.**
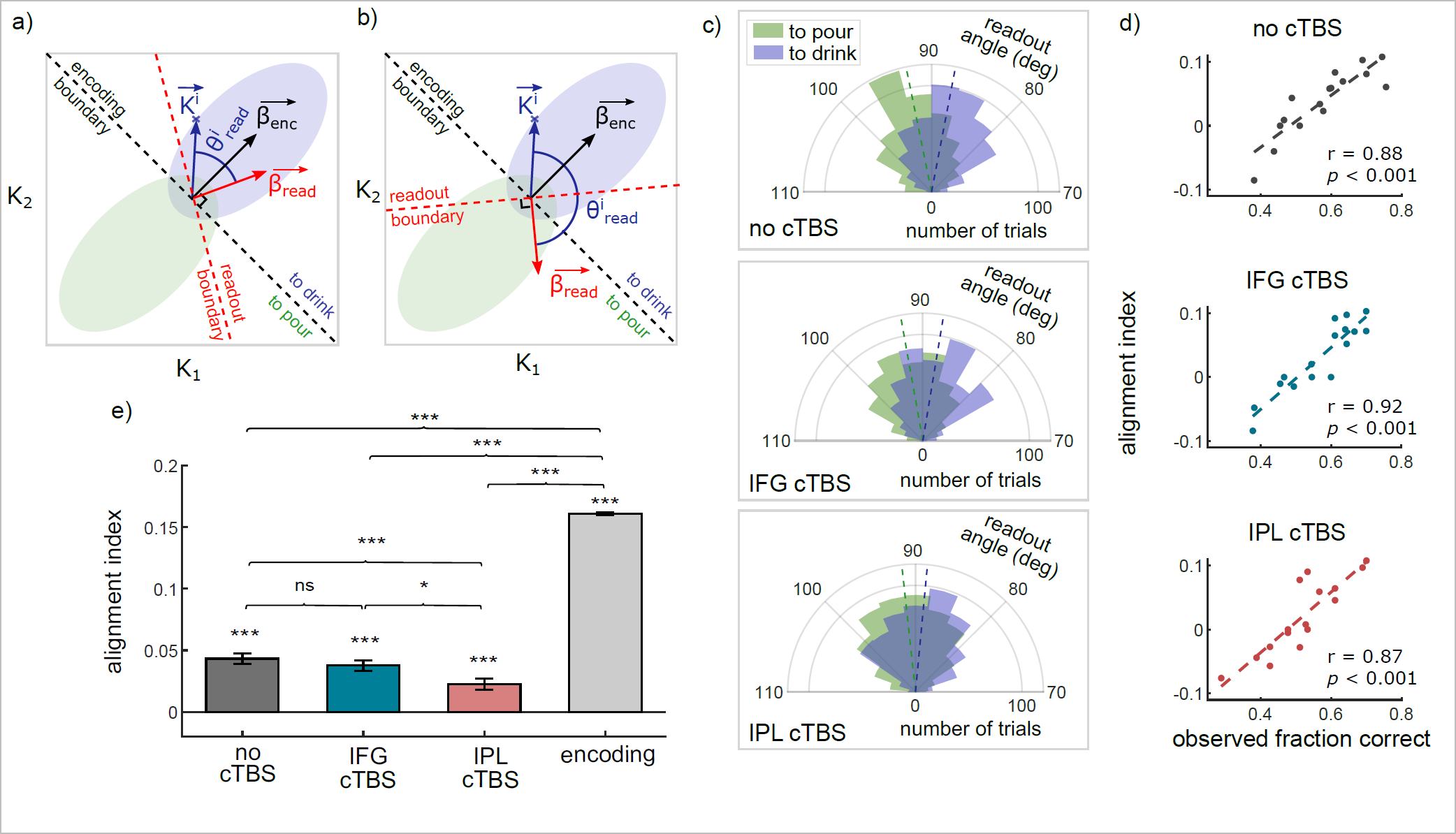
Misalignment of readout following IPL cTBS. **a-b** Diagrams illustrating alignment (a) and misalignment (b) between intention encoding and readout in a simplified two-kinematic feature space. Conventions are as in Fig. 2e and 3c. The larger the angle between the encoding vector and the readout vector, the larger on average the angle between the single-trial kinematic vector and the readout vector. This justifies an alignment index based on the cosine of the angle between the single-trial kinematic vector and the readout vector. Positive alignment indices denote correct readout. **c** Polar distribution of readout angles for ‘to pour’ and ‘to drink’ trials under no cTBS, IFG cTBS and IPL cTBS. **d** Scatterplot of alignment indices against observed task performance across participants under no cTBS, IFG cTBS and IPL cTBS. **e** Effect of IPL cTBS on the alignment index. For comparison, we show also the value of the alignment index computed as the signed cosine of the encoding angle *θ*_*enc*_. As shown in Fig 2g, the angle *θ*_*enc*_ between the encoding vector and the single-trial kinematic vector 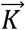 can be used to classify the intention of single-trial kinematics with 100% accuracy. Histograms represent mean ± SEM across all trials and participants.

At the single-trial level, alignment can be computed as the angle between the single-trial kinematic vector and the readout vector. Angles of 90° indicate that readout is totally unrelated to intention-information encoded in kinematics; angles approaching 0° (Fig. 4a) indicate full correct readout of intention information for ‘to drink’ trials (and totally incorrect readout of information for ‘to pour’ trials); angles approaching 180° (Fig. 4b) indicate full correct readout of intention information for ‘to pour’ trials (and totally incorrect readout of intention-information for ‘to drink’ trials). As shown in Fig. 4c, for no cTBS (and also for IFG cTBS), single-trial angle distributions were centered 3° away from orthogonality (90°), with ‘to pour’ and ‘to drink’ distributions only partly overlapping, and the majority of trials distributed in the correct readout angle range. For IPL cTBS, single-trial angles were centered only 1° away from orthogonality, with an almost complete overlap between intention-specific distributions and with about half of trials in the incorrect readout angle range (Fig. 4c). These data suggest that IPL cTBS impaired the ability to read out intention information encoded in single-trial kinematics correctly.

To quantify these observations, we devised a single-trial alignment index based on the projection of the single-trial kinematic vector on the readout vector. Although the projection is a signed valued that indicates intention, we adjusted the sign so that positive alignment indices denote correct readout and negative alignment indices denote incorrect readout (see Methods). Consistent with the observation that angle distribution was closer to 90° after IPL cTBS, results revealed a significant decrease of alignment after IPL cTBS in comparison to both no cTBS and IFG cTBS (Fig. 4e). To rule out that such a decrease could be accounted for by small (not significant) differences in model performance across sessions, we repeated the analyses considering only choices correctly predicted by the model. The pattern of results remained highly similar (Supplementary Fig. 4i-l). For the kinematic discrimination task, no cTBS and IPL cTBS did not differ in alignment (Supplementary Fig. 4a-h).

To substantiate the link between alignment and individual task performance, we quantified the fraction of behaviorally correct trials at the single-subject level as a function of alignment. For all sessions and tasks, alignment was positively correlated with individual task performance (Fig. 4d). Moreover, for both the intention and the kinematic tasks, participants who experienced a larger decrease in alignment also experienced a larger decrease in discrimination performance following IPL cTBS (Pearson’s correlation r = 0.84 for intention task, r = 0.74 for kinematic task, *p* < 0.001). Further, alignment was more predictive of individual discrimination performance than other model parameters (such as the norm and number of non-zero readout coefficients; see Supplementary Data 6 and Supplementary Fig. 5). Together, these results suggest that transient disruption of IPL selectively misaligned intention readout with respect to encoding.

### Origins of misalignment between intention encoding and readout

To understand the origins of IPL cTBS induced misalignment, we further examined the distribution and concordance in sign of readout coefficients relative to encoding coefficients. Given the lack of effect of IFG cTBS at both subject-level and trial-level, in this analysis we focused on comparing no cTBS and IPL cTBS.

We first assessed whether misalignment could result from a shift in the distribution of readout coefficients towards non-informative kinematic features, that is, whether a larger fraction of non-zero readout coefficients are assigned to non-informative individual kinematic features after IPL cTBS. Against this hypothesis, we found no difference in the average fraction (and norm) of non-zero readout coefficients assigned to informative kinematic features between no cTBS and IPL cTBS (Fig. 5a). This indicates that transient disruption of activity in IPL did not alter the relative weight given to informative versus non-informative kinematic features.

**Fig. 5.**
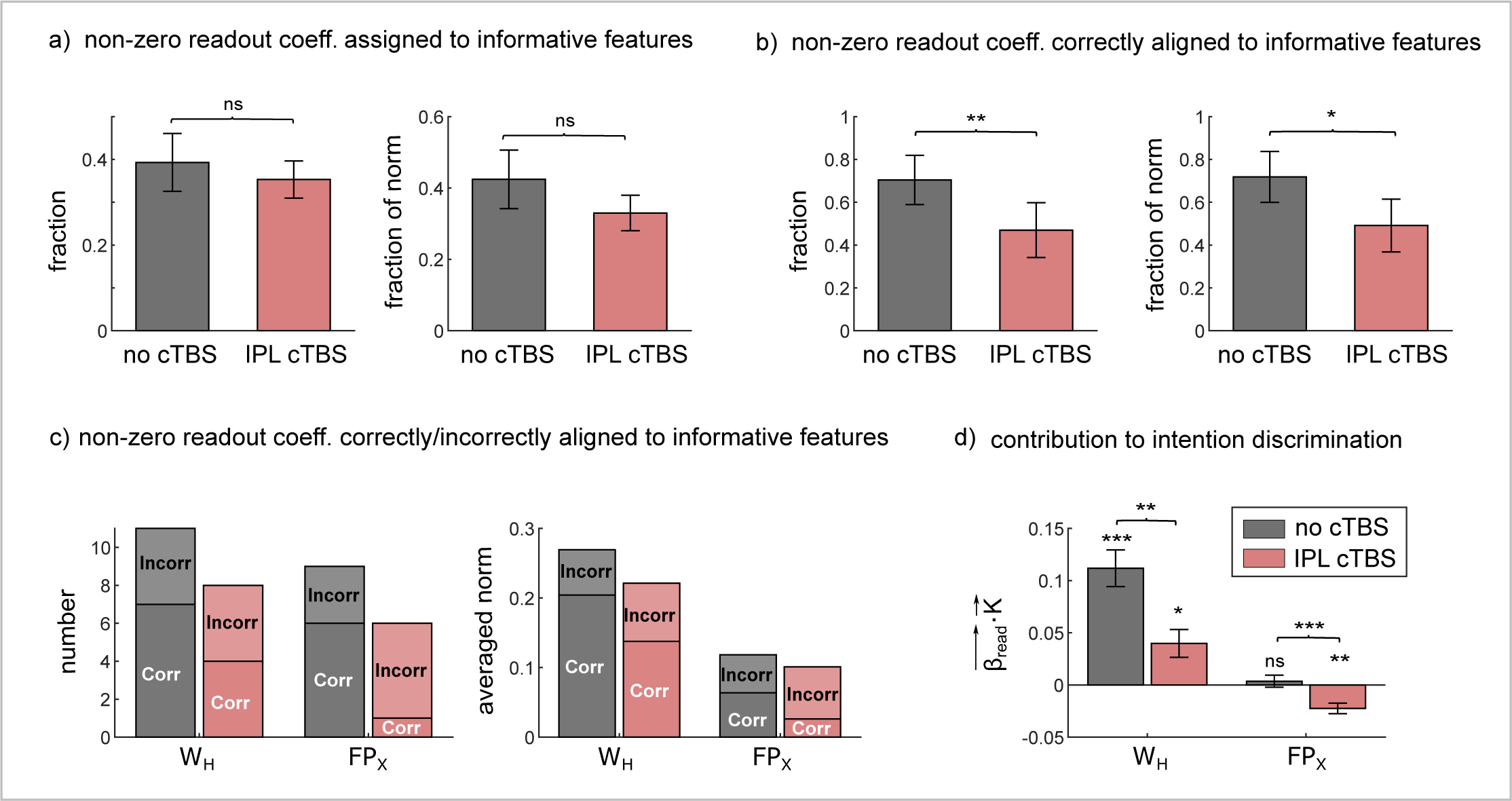
Origins of misalignment. **a** Fraction and fraction of the norm of non-zero readout coefficients assigned to informative features. **b** Fraction and fraction of the norm of non-zero readout coefficients assigned to informative features and correctly aligned with encoding. Fraction was computed on a subject basis and then averaged across subjects. **c** Number and averaged norm of non-zero readout coefficients (in)correctly aligned to informative features in encoding. We focused on the most informative and most read out kinematic variable: the height of the wrist (W_H_) and the relative abduction/adduction of the thumb and the index finger, irrespective of wrist rotation (FP_X_). **d** Contribution of W_H_ and FP_X_ to intention discrimination performance, computed as the scalar product between the kinematic vector and the readout vector within that feature subspace. Histograms represent mean ± SEM across all trials and participants.

A second possible origin of misalignment is that IPL cTBS deteriorates the mapping from informative kinematic features to intention choices. For example, a variation in a particular feature encoding ‘to drink’ (e.g., higher wrist height at 75% of movement duration; Fig. 2a), correctly read as ‘to drink’ before cTBS, might be incorrectly read as ‘to pour’ following cTBS. Geometrically, this would correspond to the readout vector and the encoding vector pointing in opposite directions (Fig. 4b), that is, the coefficients of the readout vector and of the encoding vector for the considered feature having opposite signs. Consistent with this hypothesis, the fraction (and norm) of non-zero readout coefficients correctly aligned to informative kinematic features was lower in IPL cTBS compared to no cTBS (Fig. 5b). In contrast, no difference between no cTBS and IPL cTBS was observed for the kinematic discrimination task (see Supplementary Fig. 6a, b). Together, these analyses indicate that cTBS to the left IPL altered the mapping of informative kinematic features to intention choices.

### IPL cTBS alters readout of the two most informative kinematic features

To gain further insight into the inter-individual reproducibility of readout, we explored how specific features were read by different observers by computing cross-correlations between the readout weights (computed as regression coefficients) of different participants. Observers showed significant but overall moderately low cross-correlations between readout weights (see Supplementary Data 3), attesting to a diverse range of individual readout patterns. Interestingly, under no cTBS, cross-correlation values were significantly larger for informative features than for non-informative features. This indicates that readouts of non-informative features were more variable than those of informative features. After disruption of IPL, cross-correlation values no longer differed between informative and non-informative features, implying that readout variability was no longer reduced for informative features (see Supplementary Data 3).

Strikingly, under no cTBS, the two variables that were read out more consistently across observers (see Supplementary Data 3 and Supplementary Fig. 3c) were also more informative over a wider time range in terms of encoding (Supplementary Fig. 3a): the height of the wrist (W_H_) and the relative abduction/adduction of the thumb and the index finger, irrespective of wrist rotation (FP_X_). Under no cTBS, for W_H_, readout weights correctly aligned with respect to encoding outweighed incorrectly aligned readout weights both in number and norm (Fig. 5c). This indicates that most W_H_ readouts were correctly aligned. For FP_X_, in contrast, correctly aligned readout weights were greater than incorrectly aligned ones in number but not in norm (Fig. 5c), suggesting less appropriate readouts. IPL cTBS decreased the number and the norm of correctly aligned readout weights for both kinematic variables. For FP_X_, but not for W_H_, this decrease was accompanied by an increase in number and norm of readout weights incorrectly aligned relative to encoding.

To estimate the implications of these readout patterns for single-trial discrimination performance, we computed an index of how much W_H_ and FP_X_ contributed toward correct readout of intention information and thus correct choice. This index was computed as the scalar product between the kinematic vector and the readout vector within the kinematic subspace formed by each variable. We adjusted the sign so that positive values of this index indicate positive contribution toward correct readout. As shown in Fig. 5d, whereas the contribution of W_H_ to single-trial task performance was positive under no cTBS and remained positive after IPL cTBS, the contribution of FP_X_ changed from null to negative, suggesting that, following IPL cTBS, incorrect readout of FP_X_ contributed toward decreased task performance. For the kinematic discrimination task, IPL cTBS had no influence on how the most informative variables, including W_H_, were read out (Supplementary Fig. 6c, d). These findings fit well with the above reported result of a selective decrease in alignment between intention encoding and readout after IPL cTBS, and demonstrate how such decrease directly affected the most informative and most read out kinematic variables in the intention discrimination task.

## Discussion

While theoretical and experimental work suggest that attribution of intention to action is mediated by the frontal-parietal action observation network, the specific neural computations involved in this ability have remained largely unknown and causally untested. Here we developed a novel approach, combining offline cTBS with analytical methods inspired by neural information coding, to reveal single-trial computations in the left anterior IPL and the left IFG pars orbitalis. As demonstrated by our re-analysis of fMRI data [9], both these regions predict the intention attributed to observed reach-to-grasp acts. However, whether they play a causal role in intention to action attribution was undetermined.

Our work causally establishes the role of the left IPL in intention to action attribution, which has long been hypothesized, but not directly tested [3]. We presently demonstrate that cTBS applied over a left anterior IPL site, but not over a left IFG pars orbitalis site, impairs the ability to discriminate intention from variations in reach-to-grasp kinematics. Confirming the selectivity of the reported effects to intention discrimination, IPL cTBS does not affect the ability to discriminate variations in individual kinematic features unrelated to intention (kinematic discrimination task). These findings provide evidence of a functional role of the left anterior IPL in attributing intentions to actions. But what are the computations involved in this ability? What (and how) does the left anterior IPL compute?

Our encoding results indicate that variations in kinematic features accurately specify intention information, with a very small fraction of kinematic variance affording a perfect intention encoding. In the absence of perturbation of neural activity, intention choice relies on the selective readout of such information at the single-trial level. Our combined experimental and modeling results indicate that the pattern of readout is sparse and idiosyncratic, in that individual observers read different features. Across trials, however, informative features are read more, assigned larger readout weights and read more correctly than non-informative features. Notably, we found that the two features more consistently read out across observers are also the two features informative over a wider time range in terms of encoding. Hence, observers place greater weight on features that carried more intention information. This suggest that while intention readout does not optimally exploit intention-information encoded in movement kinematics, it consistently relies on the most informative features.

Transient disruption of left anterior IPL does not impair the ability to ‘see’ changes in specific kinematic features, nor does it alter the relative weight given to informative versus non-informative features. Rather, it selectively decreases alignment between intention encoding and readout, affecting the observer’s ability to link variation in informative kinematic features to the correct intention.

These results provide causal support for an architecture in which left IPL represents goals or intentions and is placed at the highest level of the cortical hierarchy engaged during action observation [3]. Kinematics, goal-object and intention are generally conceived as independent levels of such hierarchy [36], with IPL neural populations responding to intention as opposed to kinematics. However, our finding that transient disruption of activity in left anterior IPL affects the correct readout of informative kinematic features suggests that the responsivity of IPL neurons to kinematics should be reconsidered. We speculate that, while neurons in the left anterior IPL do not code variations in individual kinematic features, they respond to intention-related variations in movement kinematics.

In this framework, it may be surprising that transient disruption of activity in the left IFG pars orbitalis did not affect intention discrimination. Both monkey and human studies relate left IFG to intention coding [6–9]. Moreover, there is evidence that in humans repetitive TMS applied to the left IFG impairs the interpretation of observed kinematic patterns [27]. The lack of behavioral modulation to IFG pars orbitalis in our study may indicate that, while potentially accessible to a classifier as demonstrated by our MVPA re-analysis of fMRI data, intention-information in the left pars orbitalis does not functionally contribute to intention discrimination. Alternatively, it is possible that other regions – within IFG or outside this region – compensate for perturbation of the left pars orbitalis. Plausible candidates include the left IFG pars opercularis and triangularis, as both these regions have been implicated in kinematic readout [8,37]. To distinguish between these hypotheses, future experiments will be needed, possibly combining our single-trial readout model framework with dual-coil TMS and fMRI-TMS approaches.

The set of analytical methods developed in the current framework could be further generalized to examine how humans come to read a variety of mental states encoded in the movements of the eyes, mouth, hands and body. Moreover, our approach could be useful for developing intuition about how atypical encoding and readout link to deficits in social cognition [38]. For example, individuals with autism have difficulties perceiving, predicting and interpreting the actions of others [39]. The analysis and methods presented here could provide a useful tool for generating and testing alternative hypotheses about how altered readout computations affect ability to make inferences about others’ mental states.

## Author Contributions

C.B. and S.P. conceived and supervised the project. C.B. and A.C. designed the experiments. S.P., C.B. and J-F.P. designed the analyses, with contributions from M.V. M.S. and A.K. performed the experiments. J-F.P., K.P. and A.C. performed the analyses, with contributions from M.V. A.A. advised on cTBS. C.B., S.P. and J-F.P. wrote the manuscript. All authors contributed to the content and writing of the Supplementary Information.

## STAR Methods

### Resources Availability

Further information and requests for resources should be directed and will be fulfilled by the Lead contact, Cristina Becchio.

### Experimental model and subject details

Based on [40], we decided *a priori* to collect data from at least 15 participants in each task. To achieve this, we had 20 participants perform the intention task and 20 participants perform the kinematic discrimination task. Three participants were removed from the sample as they did not complete all the three sessions. Additionally, two participants were unable to complete the cTBS sessions due to a too-high resting motor threshold (above 80% of maximal stimulator output). Thus, *N =* 16 for the intention discrimination task (10 females, 6 males, mean age 23, range 19-27 years) and *N =* 19 for the kinematic discrimination task (9 females, 10 males, mean age 24, range 20-28 years). All participants were right-handed according to the Edinburgh Handedness Inventory [41] and had normal or corrected to normal vision. None of the participants reported neurological, psychiatric, or other medical problems or any contraindication to MRI or TMS [42,43]. Informed written consent was obtained in accordance with the principles of the revised Helsinki Declaration (World Medical Association General Assembly, 2008) and with procedures cleared by the local ethics committee (Comitato di Bioetica di Ateneo, University of Turin). All participants received monetary compensation for their time.

### Method details

#### Experimental design and procedures

The design of the intention discrimination and kinematic discrimination tasks was between-subjects, while effects of cTBS used a within-subject design. Participants assigned to each task underwent a high-resolution MRI structural scan, after which they attended three experimental sessions: no cTBS, cTBS to the left IPL and cTBS to the left IFG. During each of these sessions, participants completed the intention discrimination task (or the kinematic discrimination task, depending on task assignment) followed by a control contrast discrimination task. Participants sat in front of a 24-in. inch computer screen (resolution 1280 x 800 pixels, refresh frequency 60 Hz) at a distance of 50 cm in a dimly lit room. Each session lasted approximately 90 minutes and occurred at the same time of the day (±1 h) for each participant. Participant sessions were separated by one week, and session type order was randomized across participants.

#### MRI acquisition

T1-weighted scans were acquired using a 1.5 Tesla INTERA™ scanner (Philips Medical Systems) equipped with a 32-channel SENSE high-field head coil. Each high-resolution structural scan included 160 axial slices with an in-plane field of view (FOV) of 256 × 240 and a gap of 0 mm for a resolution of 1 × 1 × 1 mm (TR = 8.2 ms, TE = 3.80 ms, flip angle = 8 degrees). T1-weighted scans were used for the MRI-guided neuronavigation used to target cTBS stimulation sites (see below).

#### MRI-guided cTBS protocol

MRI-guided cTBS was administered using a 70-mm figure-eight coil connected to a Magstim Rapid2 stimulator (Magstim, Dyfed, UK). A SofTaxic NeuroNavigator system (EMS, Bologna, Italy) was employed to determine the coil position for all the ‘to-be-stimulated’ brain regions. Specifically, individual MRI scans were used to first construct scalp surface and skull landmarks of the left periauricular (A1), right periauricular (A2) and nasion (N) on the participant’s T1 MRI image. The brain scan was then normalized to Talaraich space and neuronavigation data were co-registered to measurements taken from the same points of reference (A1, A2, N) sampled from the participant’s scalp. The intensity for the cTBS protocol was set at 90% of the resting Motor Threshold (rMT), defined as minimal stimulation intensity producing motor evoked potentials (MEPs) of a minimum amplitude of 50 µV in the first dorsal interosseous (FDI) muscle [42]. To determine the rMT, for each participant for each stimulation session, we applied single pulse TMS over the left Primary Motor Cortex (M1) and recorded the MEPs from the right FDI muscle using a Biopac MP-150 (Biopac Systems, Inc., Santa Barbara, CA) through pairs of Ag–AgCl surface electrodes in a belly tendon montage. The rMT was determined by means of adaptive parameter estimation by sequential testing procedure (PEST) with the Motor Threshold Assessment Tool 2.0 [44]. The coordinates for targeting left M1 (tal x = -44, y = -19, z = 53) were extracted from the neurosynth reverse inference map for the term ‘index finger’ [45]. Following the rMT estimation procedure, cTBS was delivered to the left IFG and the left IPL targets. cTBS consisted of three pulses at 50 Hz repeatedly applied at intervals of 200 ms (5 Hz) for 40 s [46]. In cTBS sessions, discrimination tasks were administered 5 min post cTBS, that is, in the time window in which maximal inhibitory effects of stimulation have been reported [46–50].

#### cTBS targets

cTBS targets were defined based on a re-analysis of fMRI data published in [9]. Using the same set of action stimuli used in the current cTBS experiments, [9] found that the left IPL and the left IFG represent, among other areas, the intention of an observed reach-to-grasp act performed with the intent to pour or to drink. Here, we used multi-voxel pattern analysis (MVPA) to identify intention-sensitive targets that could accurately discriminate intention within these regions. First, we defined separate anatomical regions of interest (ROI) using the Automated Anatomical Labeling (AAL) [51] library contained in MarsBaR SPM toolbox [52]. We extracted separate ROIs for the left IFG pars opercularis, pars triangularis, and pars orbitalis, and the left IPL. Next, for each ROI, we trained and tested separate linear SVM classifiers to distinguish between the two intentions with accuracy assessed using leave-one-subject-out (LOSO) cross validation. For the left IFG, intentions were classified most accurately in the pars orbitalis (classification accuracy = 0.73; *p* < .01, permutation test). The MVPA-defined cTBS target (MNI x = -46, y = 30, z = -12, then converted to Talairach stereotactic coordinates x = -46, y = 29, z = -12) was chosen in a portion of the pars orbitalis that contained a large number of informative voxels (20% highest ranked voxels; see Supplementary Fig. 7). For left IPL, intentions were classified with an accuracy of 0.78 (*p* < .001, permutation test). The MVPA-defined cTBS target (MNI coordinates x = -58, y = -34, z = 40, then converted to Talairach stereotactic coordinates x = -57, y = -31, z = 38) was chosen in the anterior part of the IPL [53], which again contained a large number of voxels ranked as most informative (20% highest ranked voxels; Supplementary Fig. 1a). cTBS targets were identified for each participant on their T1-weighted MR-scan. The SofTaxic neuronavigational system (E.M.S. srl, Bologna, Italy) located the corresponding scalp position with an error threshold set to the default value of 2 mm.

#### Action stimuli

Stimuli were selected from a dataset of 512 grasping acts obtained by recording 17 naïve participants reaching toward and grasping a bottle with the intent to pour some water into a small glass or to drink water from the bottle. Detailed apparatus and procedures are described in [40]. Briefly, reach-to-grasp movements were tracked using a near-infrared camera motion capture system with nine cameras (frame rate, 100 Hz; Vicon System) and concurrently filmed using a digital video camera (Sony Handy Cam 3-D, 25 frames/sec). Sixteen kinematic variables of interest were computed throughout the reach-to-grasp phase of the movement, from reach onset to reach offset [54]. A list of variables and how they are computed is reported in Supplementary Table 1 and in Supplementary Methods 1. Sixty grasping acts (grasp-to-pour, *N* = 30; grasp-to-drink, *N* = 30) were selected to satisfy the following requirements: i) within-intention distance was minimized (using the metric reported in [40]); ii) median split based on maximum wrist height led to a significant difference between “higher” and “lower” wrist height grasps (t_58_ = 11.2; *p* < 0.001); iii) maximum wrist height did not differ between intentions (*p* = 0.27). The corresponding movies, filmed from a lateral viewpoint, were used as stimuli in the intention discrimination task and in the kinematic discrimination task. Movies were edited with Adobe Premiere Pro CS6 (mp4 format, disabled audio, 25 frames per second, resolution 1,280 × 800 pixels) so that each movie clip started with the reach onset and ended at contact time between the hand and the bottle. Movement duration (mean ± SEM = 1.04 ± 0.02 s, range = 0.84 to 1.36 s) did not differ between intentions (t_58_ = -0.30; *p* = 0.76).

#### Intention discrimination task

The intention discrimination task consisted of two blocks of 60 trials. Task structure conformed to a 2AFC design. Each trial displayed two reach-to-grasp acts in two consecutive temporal intervals: one interval contained a grasp-to-pour act, the other a grasp-to-drink act. Depending on block, participants had to indicate the interval (first or second) containing the grasp-to-drink or grasp-to-pour act. Each trial started with the presentation of a white central fixation cross for 1500 ms. Then, the first grasping act was presented followed by an inter-stimulus interval of 500 ms, after which the second grasping act was presented. After the end of the second video, the screen prompted participants to indicate the interval (first or second) containing the grasp-to-drink (or grasp-to-pour, depending on block) action by pressing a key. The prompt screen was displayed until response or for a maximum duration of 3000 ms. After response, participants were requested to rate the confidence of their choice on a four-level scale by pressing a key. Pairing of videos was randomized across trials and participants. To ensure that grasping actions could be temporally attended (i.e., to allow participants enough time to focus on movement start), 9, 11, or 13 static frames were randomly added at the beginning of each video. In order to equate video durations, static frames were also added at the end of each videos in a compensatory manner. Participants began the session by performing a practice block before the main experimental task. The order of the presentation of the blocks was counterbalanced across participants. Stimulus presentation, timing and randomization was controlled using E-prime V2.0 software (Psychology Software Tools, Pittsburgh, PA).

#### Kinematic discrimination task

The kinematic discrimination task included the same stimuli and design as the intention discrimination task, except that participant were asked to indicate the interval containing the grasp with higher (or lower, depending on block) peak vertical height of the wrist.

#### Control contrast discrimination task

To control for cTBS effects unrelated to action observation, such as integration of evidence favoring one alternative over time, participants performed a contrast discrimination task at the end of each session. The contrast discrimination task consisted of three blocks of 32 trials. Each trial started with the presentation of a fixation cross (1000 ms), after which two grey rectangles were displayed for 1000 ms on two consecutive intervals separated by a 500 ms inter-stimulus interval. In half of the trials, the rectangles had the same contrast (rgb = 100,100,100). In the other half of the trials, the difference in contrast was of 10, 15, 20 or 25 in the rgb space. For each trial, the participant had to indicate whether the contrast of the rectangles was ‘same’ or ‘different’ (within a 3000 ms window) and rate the confidence of their choice on a four-level scale by pressing a key. Results are reported in Supplementary Data 7.

### Data Analyses

#### Data preprocessing

Trials for which subjects failed to provide a response within 3000 ms were discarded from the analyses (0.5% of trials for the intention discrimination task and 0.1% of trials for the kinematic discrimination task). The first 25% of trials in each block were discarded to account for the time needed by participants to familiarize with task and response mapping. As a control, we verified that the pattern of results and their significance remained similar even when including all trials. Specifically, similarly to Fig. 1d, e, we found a significant decrease in intention discrimination post IPL cTBS relative to both no cTBS and IFG cTBS. Kinematic discrimination did not differ across sessions.

#### Single-trial kinematic vector

To model single-trial kinematics, we first averaged, for each grasping act, the 16 kinematic variables of interest over four epochs of 25% of the normalized movement time (0-25%, 25-50%, 50-75%, and 75-100% of movement duration defined from reach onset to reach offset). Next, for each trial, we combined the kinematic features associated with the two grasping acts in a 64-dimensional kinematic vector, 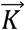, defined as:

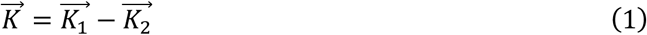

where 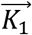 and 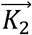 denote the vectors of kinematic features associated to the first and second reach-to-grasp act displayed in the trials. This definition, used in all of our logistic regression analyses, reflects the assumption that, in a 2AFC task, choices are based on comparative judgements. Using more detailed regression models that employed the kinematic features of the two grasping acts did not improve model predictability (see Supplementary Data 2).

#### Logistic regression models of encoding and readout

We analyzed encoding and readout using two sets of logistic regression models: encoding and readout models. Logistic regression [35] is a linear regression for log-odds, and is a standard probabilistic approach to classification. Logistic regression models are powerful for explaining behavioral strategies [55]. They confer several advantages in modeling the dependence of a random binary variable, such as observers’ choice, on one or more explanatory variables [56]. For example, they assume binomial noise – the most natural noise model for binary responses; they combine predictor variables linearly; they can be robustly fit to data; they have a graded nonlinearity, which allows for a modulation of probabilities different from an all-or-none binarization. The latter property was particularly suitable to the readout model because in our data discrimination performance was positively correlated with confidence ratings in the no cTBS session (Spearman’s correlation: *p* < 0.05 and *p* < 0.001 for the intention and kinematic discrimination tasks, respectively), suggesting a graded nature of the response probability as a function of the kinematic evidence. To aid comparison between encoding and readout models, we also used logistic regression for modeling encoding. Versions of the encoding models based on other formulations, such as linear discriminant analyses, were also built and tested, and yielded qualitatively similar results.

The logistic model expressed the probability of a binary stochastic variable *Y*, where *Y* takes the values ‘to drink’ and ‘to pour’ for intention discrimination, as a sigmoid transformation of the sum of the components of the single-trial kinematic vector 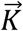. The equation of the model was as follows:

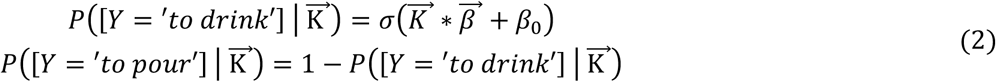

where *σ* is the sigmoid function, 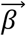 is the vector containing the values of the regression coefficients of each kinematic feature, and *β*_*0*_ is the bias, kinematic independent, term.

#### Training logistic regression models

Training and evaluation were performed similarly for both sets of models and for both discrimination tasks. Each model was trained on the set of the 90 trials retained for analyses. We z-scored the single-trial kinematic vectors within each model in order to avoid penalizing predictors with larger ranges of values. To avoid over-fitting, we trained each model using elastic-net regularization, with a value of *α* = 0.95 for the elastic net parameter, which provides sparser solutions in parameter space [57]. We verified that our results were robust to the choice of elastic net regularization (see Supplementary Data 3). The free parameter λ, which controls the strength of the regularization term, was estimated for each model using leave-one-out cross-validation. We retained for each model the value λ_min_ associated to the minimum mean cross-validated error. Models were then trained on all 90 trials with the retained regularization term. Logistic regression was implemented using *R glmnet* package [58]. In the main text, we report the results obtained by applying this training procedure as it gives only one set of regression coefficients per analyzed case and it is therefore easier to interpret. However, qualitatively similar results were obtained when using leave-one out cross validation on the entire procedure (on top of the cross validation used for the determination of the λ parameter; see Supplementary Data 3).

The following sections describe encoding and readout models with reference to the intention discrimination task. The procedures for the kinematic discrimination task were identical.

#### Encoding model

The encoding model expressed the probability of the grasping act displayed in the first interval of a given trial being ‘to drink’ as a function of the kinematic vector measured in the same trial. Having verified that intention information slightly varied as a function of video pairings (which was randomized across trials and participants), we trained the encoding model separately on each set of video pairings presented in each session to each observer. We used the encoding model to evaluate the overall amount of intention information in movement kinematics (Fig. 2g, Supplementary Fig. 3a, b).

#### Readout model

The readout model expressed the probability of intention choice in a given trial as a function of the kinematic vector measured in the same trial. We trained the readout model separately for each observer in each session. To model intention choice as a function of single-trial kinematics, we trained the readout model using all 64 kinematic features (16 kinematic variables at four time epochs).

#### Evaluation of model performance

To quantify model performance (Fig. 2g, 3d,g), we computed for each trial the most likely value of the variable *Y* by taking the argmax over *Y* of 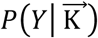 in Eq. 2. This parameter provides an estimate of the model prediction (prediction of the actual intention for the encoding model; prediction of observer’s choice for the readout model) on a single trial. We then quantified model performance as the fraction of correct predictions computed over all the trials. The chance-level null-hypothesis distribution for readout model performance was created by fitting the model after randomly permuting across trials the observer’s choice labels.

#### Computation of task performance predicted by the readout model

In Fig. 3e,h, we used the readout model to estimate, for each participant, the fraction of behaviorally correct trials. Using the logistic readout model (Eq. 2), we computed for each trial the probability of each choice. Then we averaged across all trials the probability of the correct choice.

#### Classification of individual kinematic features as informative

To evaluate the informativeness of individual features about intention (which is used for Fig. 5, Supplementary Fig. 3a, b, and Supplementary Fig. 6), we used a single-feature encoding model implemented using Matlab’s *glmfit* function. Each model was trained on the full set of 90 trials retained for analyses. We z-scored the single-trial kinematic vectors. Significance of the regression coefficient was assessed with t-statistics. We retained as informative kinematic features whose regression coefficients were found to be significant (*p* < 0.05) in all video pairings. In Supplementary Fig. 3a, b, variables are ranked in terms of their encoded information. Ranking was determined by computing the average norm of the regression coefficient of each informative feature across all video pairings and then summing the obtained norms across features belonging to the same kinematic variable.

#### Contribution of individual kinematic variables to discrimination performance

Fig. 5d and Supplementary Fig. 6d visualize the contribution of individual kinematic variables to discrimination performance. Single variable contribution to discrimination performance was computed as the scalar product between the kinematic vector and the readout vector calculated within the feature subspace formed by the features of the considered kinematic variable (e.g., 25%, 50%, 75% and 100% of movement time for W_H_). Positive values of this index imply a positive contribution of the variable towards enhancing discrimination performance; negative values imply a negative contribution towards decreasing discrimination performance.

### Quantification and Statistical Analysis

#### Short summary

The *p* values of all reported statistical comparisons are two-sided and Holm-Bonferroni corrected. Details are reported below.

#### Conventions for plotting p values

In all figures, * indicates *p* < 0.05, ** indicates *p* < 0.01, and *** indicates *p* < 0.001, *ns* indicates *p > 0.05*. Following standard notations, * above bars indicate significance of difference from chance of an individual quantity, * above brackets indicate significance of difference between two quantities.

#### Logistic Mixed Effects Models for assessing differences in the distribution of binary variables

We assessed the significance of differences in discrimination performance (Fig. 1d, e), response bias (Supplementary Fig. 1b), and readout model performance (Fig. 3d, g) against chance and across sessions using Logistic Mixed Effects Models (LMEM) (see Supplementary Data 1). Logistic statistics were used because these quantities are computed from binary stochastic variables in each trial and thus cannot be assessed with t-tests or other parametric Gaussian statistics. The logistic regression estimated the value and standard error of each effect, from which a *z* value (for computation of two-sided *p* values) and a Cohen’s *d* value (for estimation of effect size) were computed. All *p* values were Holm-Bonferroni corrected for three comparisons (IPL cTBS vs. no cTBS, IFG cTBS vs. no cTBS, and IPL cTBS vs. IFG cTBS).

#### Permutation test for assessing differences in readout coefficients and alignment

For assessing differences in alignment (Fig. 4e, Supplementary Fig. 4d-f) and in the contribution of individual features to discrimination performance (Fig. 5d and Supplementary Fig. 6d), we used non-parametric permutation statistics based on constructing a null-hypothesis distribution of differences in values after randomly permuting session labels across trials. For assessing significance of differences in the readout regression coefficients computed for each subject across sessions (Fig. 5a, b and Supplementary Fig. 6a, b), we used a similar session label permutation test for regression coefficients. For all tests, the null-hypothesis distribution was computed using 10^4^ random permutations. In all permutation tests, reported *p* values are two-sided.

#### Significance of correlations

Significance of Pearson’s correlation values and step-wise regression coefficients were assessed using two-sided parametric Student statistics [59] implemented in the MATLAB functions *corr* and *stepwisefit*, respectively. We assessed significance of Spearman correlations using the two-sided permutation distribution [60] implemented in the MATLAB function *corr*.

### Data and Code Availability

Data are available upon request to the Lead contact, Cristina Becchio. Custom MATLAB code for encoding and readout single-trial analyses will be made publicly available on Github before publication.

## Supplementary Information

### Supplementary Methods

***Supplementary Methods 1.*** *Computation of kinematic variables*

We used a custom software (Matlab; MathWorks Inc., Natick, MA) to compute two sets of parameters of interest: F_global_ and F_local_ parameters. F_global_ parameters were expressed with respect to the global frame of reference, i.e., the frame of reference of the motion capture system. Within this frame of reference, we computed the following parameters:

a. Wrist Velocity, defined as the module of the velocity of the wrist marker (mm/sec);
b. Wrist Height, defined as the z-component of the wrist marker (mm);
c. Wrist Horizontal Trajectory, defined as the x-component of the wrist marker (mm);
d. Grip Aperture, defined as the distance between the marker placed on thumb tip and the one placed on the tip of the index finger (mm). To provide a better characterization of the hand joint movements, the second set of parameters was expressed with respect to a local frame of reference centered on the hand (i.e., F_local_). Within F_local_ we computed the following parameters:
e. x-, y-, and z-thumb defined as x-, y- and z-coordinates for the thumb with respect to F_local_ (mm);
f. x-, y-, and z-index defined as x-, y- and z-coordinates for the index with respect to F_local_ (mm);
g. x-, y-, and z-finger plane defined as x-, y- and z-components of the thumb-index plane, i.e., the three-dimensional components of the vector that is orthogonal to the plane, providing information about the abduction/adduction movement of the thumb and index finger irrespective of the effects of wrist rotation and of finger flexion/extension;
h. x-, y-, and z-dorsum plane defined as x-, y- and z-components of the radius-phalanx plane, providing information about the abduction, adduction and rotation of the hand dorsum irrespective of the effects of wrist rotation.

**Supplementary Table 1.**
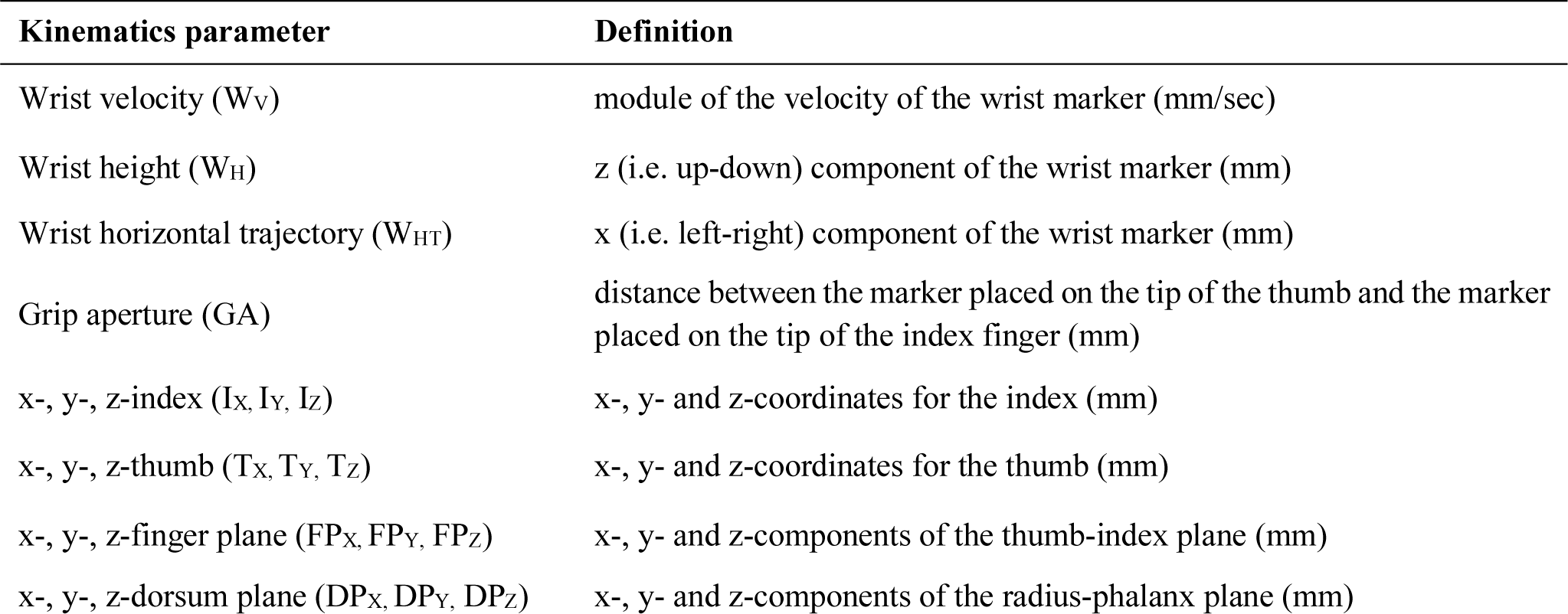
Kinematics parameters of interest. Kinematic variables were computed throughout the reach-to-grasp phase of grasp-to-pour and grasp-to-drink acts, from reach onset to reach offset

| Kinematics parameter | Definition |
| --- | --- |
| Wrist velocity ($W_v$) | module of the velocity of the wrist marker (mm/sec) |
| Wrist height ($W_H$) | z (i.e. up-down) component of the wrist marker (mm) |
| Wrist horizontal trajectory ($W_{HT}$) | x (i.e. left-right) component of the wrist marker (mm) |
| Grip aperture (GA) | distance between the marker placed on the tip of the thumb and the marker placed on the tip of the index finger (mm) |
| x-, y-, z-index ($I_x, I_y, I_z$) | x-, y- and z-coordinates for the index (mm) |
| x-, y-, z-thumb ($T_x, T_y, T_z$) | x-, y- and z-coordinates for the thumb (mm) |
| x-, y-, z-finger plane ($FP_x, FP_y, FP_z$) | x-, y- and z-components of the thumb-index plane (mm) |
| x-, y-, z-dorsum plane ($DP_x, DP_y, DP_z$) | x-, y- and z-components of the radius-phalanx plane (mm) |

### Supplementary Data

***Supplementary Data 1.*** *Details of the Logistic Mixed Effects Model (LMEM) statistics*

We performed statistical analyses of discrimination performance and response bias using Logistic Mixed Effects Models (LMEM) [1]. We considered the 0/1 variable in each trial denoting incorrect or correct response as dependent variable, session as categorical predictor and subject identity as random effect. Models differed with regard to the random effects structure. We selected the random-effect structure of the model by comparing a random intercept only model (DF 4) with a model including both random intercept and random slope (DF 9).We performed model selection using the Bayesian Information Criterion (BIC) [2], which rewards model fit and penalizes model complexity (number of DF). We carried out model fitting using the R package lme4 [3]. Significance of comparisons across sessions and against chance was assessed as significance of fixed effects using two-tailed Wald statistics. The *p* values were Holm-Bonferroni corrected for three comparisons (no cTBS vs. IPL cTBS, no cTBS vs. IFG cTBS, and IPL cTBS vs. IFG cTBS). We used the same LMEM (with 0/1 variable in each trial denoting incorrect/correct response predicted by the model) for assessing changes in readout model performance across sessions. Details of performance of the LMEMs tested for selection are reported in Supplementary Table 2.

**Supplementary Table 2.** Comparison of the LMEMs tested for the selection of model’s random-effect structure.

| Intention discrimination task |  | Kinematic discrimination task |  |
| --- | --- | --- | --- |
| *Discrimination performance* | *BIC* | *Discrimination performance* | *BIC* |
| Random intercept*<br>*converged correctly* | 355.540 | Random intercept*<br>*converged correctly* | 393.939 |
| Random intercept and slope<br>*converged correctly* | 366.601 | Random intercept and slope<br>*boundary (singular) fit* | 406.100 |
| *Discrimination performance 10% trial selection* | *BIC* | *Discrimination performance 10% trial selection* | *BIC* |
| Random intercept*<br>*converged correctly* | 215.415 | Random intercept*<br>*converged correctly* | 206.278 |
| Random intercept and slope<br>*unable to evaluate scaled gradient* | 226.079 | Random intercept and slope<br>*failed to converge with $\max\vert \text{grad}\vert = 0.00494831$ (tol = 0.001, component 1)* | 225.392 |
| *Response Bias* | *BIC* | *Response Bias* | *BIC* |
| Random intercept*<br>*converged correctly* | 355.540 | Random intercept*<br>*converged correctly* | 393.939 |
| Random intercept and slope<br>*converged correctly* | 366.601 | Random intercept and slope<br>*boundary (singular) fit* | 406.100 |
| *Contrast Discrimination* | *BIC* | *Contrast Discrimination* | *BIC* |
| Random intercept*<br>*converged correctly* | 306.243 | Random intercept*<br>*converged correctly* | 376.575 |
| Random intercept and slope<br>*boundary (singular) fit* | 325.295 | Random intercept and slope<br>*failed to converge with $\max\vert \text{grad}\vert = 0.00608542$ (tol = 0.001, component 1)* | 393.328 |

| *Readout Model Performance* | *BIC* | *Readout Model Performance* | *BIC* |
| --- | --- | --- | --- |
| Random intercept<br>*converged correctly* | 435.040 | Random intercept*<br>*converged correctly* | 371.929 |
| Random intercept and slope*<br>*converged correctly* | 383.088 | Random intercept and slope<br>*converged correctly* | 379.901 |

\* *retained model*

***Supplementary Data 2.*** *Systematic comparison of readout models that use different numbers of kinematic features in different ways*

In all encoding and readout models presented in the main text, the single trial kinematic vector was constructed as the differences between the kinematic features of the reach-to-grasp acts displayed in the first and second interval of each trial 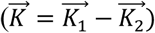.

However, we assessed whether a more complex readout model, using the full set of kinematic variables of the first and second grasping act independently (that is, 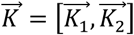), rather than their difference, would achieve better performance. The same regularization procedure, described in Methods in the main text, was applied. Performance of the model using the full set of kinematic features was not significantly different from that of the model using the difference in kinematics in any condition of the intention discrimination task (*p* > 0.4). For the kinematic discrimination task, the model using the full set of features showed a small advantage for no cTBS and IPL cTBS sessions (model performance as fraction correct: 0.88 vs 0.91, p = 0.012 for no cTBS, 0.88 vs 0.90 p = 0.044 for IPL cTBS). Both approaches achieved similarly high correlations between predicted and observed task performance (*p* < 0.001 in all cases; Supplementary Fig. 2). Given that the model using the full set of kinematic features had twice as many predictors as the model using the difference (128 vs 64 dimensions), but only led to a null to marginal increase in model performance, for the sake of parsimony, we used the simpler model in all analyses.

We also explored whether the number of time-bins used for the discretization of the temporal evolution of kinematic variables influenced model performance. We compared the performance of the four-time-bin readout model, used in all our analyses presented in the main text and supplemental figures, with that of models based on a finer time discretization (six or eight time bins). Six- and eight-time bin models performed no better than the four-time bin model (*p* > 0.4).

***Supplementary Data 3.*** *Similarity of readout weights across observers in different conditions and with different model regularizations*

To gain further insight into the robustness and inter-individual reproducibility of the readout regression coefficients, we computed cross-correlations between the readout weights of different participants under different control analyses.

We first computed the cross-correlation between the readout weights for each different pairs of participants. We found that participants showed overall moderate to low cross-correlations between their readout weights under no cTBS in the intention discrimination task (mean ± SEM across all pairs of subjects was 0.06 ± 0.02). To investigate whether the small correlation values were due to the regularization, we repeated the above analysis using different values of the hyper-parameter *α* (the one bridging continuously between Ridge regularization for *α* close to zero and Lasso regularization for *α* close to one) ranging from 0.5 to 1. This range of elastic net hyper-parameter *α* covers the range of regularization further away from the Ridge and closer to the Lasso one, which is the most suitable range given our number of trials and predictors. Cross-subject correlations slightly increased with lower *α* values, but remained always below 0.1. Similar results were obtained for IPL cTBS and IFG cTBS. As expected by design, cross-correlations in the kinematic discrimination task were much higher (mean ± SEM across all pairs of subjects was 0.39 ± 0.02 in no cTBS session), and remained high for all *α* values. These analyses corroborate the idea that the small correlation values in the intention discrimination task reflected genuine cross-subject differences.

We also considered whether cross-correlation of readout patterns differed between non-informative and informative kinematic features in the intention discrimination task. We considered separately the 19 out of 64 features that carried significant intention-related information (*p* < 0.05) (see Methods, *Classification of individual kinematic features as informative*) and computed the cross-subject correlation value in this subset of variables. Next, we compared this value with that obtained when considering the 19 non-informative features with the smallest encoding readout coefficients. Under no cTBS, cross-correlation values were significantly larger for informative features than for non-informative features (*p* < 0.05). After IPL cTBS, cross-correlation values did not differ (*p* = 0.14) between informative and non-informative features. In the kinematic discrimination task, cross-correlations values were significantly larger for informative features than for non-informative features in all conditions (*p* < 0.001).

To check whether the pattern of readout weights was robust to the choice of the elastic net hyper-parameter *α*, we computed for each participant the correlation between the readout weights for *α* = 0.95 with the readout weights obtained with values of *α* ranging from 0.5 to 1. Correlation values decreased with *α* values but remained higher than 0.9 for all *α* > 0.5. Altogether, these results indicate that readout results were robust to the choice of hyper-parameter *α*.

***Supplementary Data 4.*** *Fully cross-validated model performance and cross-validated alignment results*

In the analyses reported in the main text, although model regularization hyper-parameters were determined on the training set only, model performance was evaluated on the same set of trials used for training. Results of this analysis are thus only partly cross-validated. To control for residual overfitting, we recomputed all our main analyses using a different partitioning of data into training and testing (leave-one-out cross-validation). The pattern of results obtained with this cross-validated approach was consistent with that reported in the main text. In particular, the fully cross-validated performance of encoding models remained close to 100% for both the intention and the kinematic discrimination tasks (99.5±0.1% and 98.1±0.2% for the intention discrimination task and kinematic discrimination task, respectively). The fully cross-validated performance of readout models also remained significantly above chance in all sessions in both tasks (p < 0.001, adjusted for the three comparisons in each task). The correlation between observed and predicted task performance also remained significant in all cases (*p* < 0.001). Moreover, alignment was still significantly decreased after IPL cTBS as compared to no cTBS (*p* < 0.01).

***Supplementary Data 5.*** *Verification of the statistical significance of non-zero regression weights*

To check that that the regularization was working well and that the regression coefficients with non-zero value were meaningful [4], we did a permutation test in which a null hypothesis distribution of regression weights was obtained after random permutations of the trial labels. We took the absolute value of each individual regression coefficient obtained in the permuted dataset to build a distribution of absolute values of regression coefficient expected under the null-hypothesis of no relationship between the kinematics and the variable *Y*. We verified that all non-zero beta coefficients had an absolute value that exceeded the 95th percentile of this null-hypothesis distribution.

***Supplementary Data 6.*** *Alignment is the main predictor of task performance*

To establish whether alignment was a predictor of task performance, we computed, for each participant, the Pearson correlation between the fraction of behaviorally correct trials and the alignment index. Alignment indices for participants with null readout vectors were set to zero as in this case no information can be read out. Results, plotted in Fig. 4f and Supplementary Fig. 5, showed that the alignment index was highly correlated with task performance (Pearson correlation = 0.89, p < 0.001 for the no cTBS condition). Model parameters related to the strength of readout, but not to alignment, namely, the norm of readout vector and the number of non-zero readout regression coefficients, were also considered (Supplementary Fig. 5). These parameters showed a much weaker, borderline significant, correlation with task performance under no cTBS and no correlation after IPL cTBS in the intention discrimination task. Comparable results were found in the kinematic discrimination task (Supplementary Fig. 5).

To rank the importance of the above considered model parameters in predicting changes in task performance across conditions, we conducted stepwise linear regression analyses of the log of the ratios between IPL cTBS and no cTBS task performance of each individual participant. In both the intention discrimination task and in the kinematic discrimination tasks, the alignment index was ranked as the most important predictor (Supplementary Table 3). Only in the kinematic discrimination task, the norm of the readout vector added a significant contribution as second predictor (*p =* 0.003). The number of non-zero readout regression coefficients never added predictive power (*p >* 0.05). Collectively, these analyses support the conclusion that the alignment index was the main predictor of the change in task performance between no cTBS and IPL cTBS for each individual participant.

**Supplementary Table 3.**
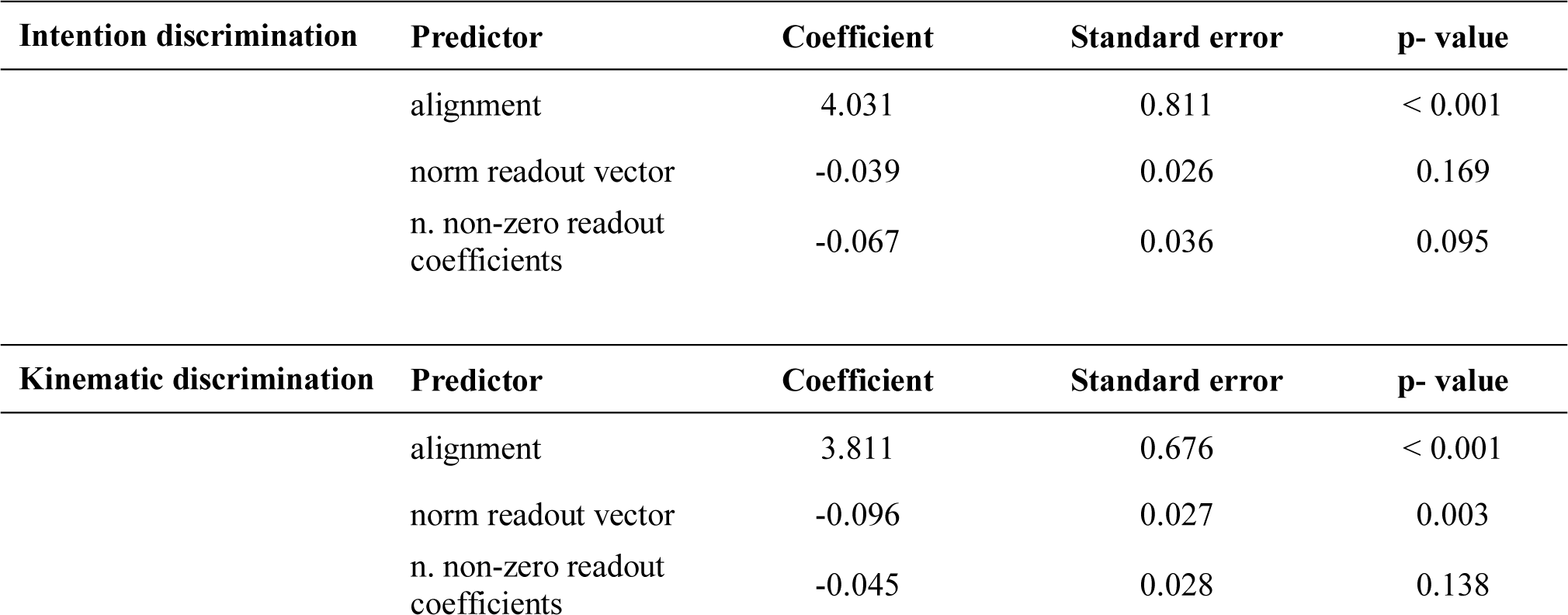
Results of stepwise regression analyses to determine the relative importance of different model parameters for discrimination performance. Predictors are ordered and listed from top to bottom, in terms of importance as computed by the stepwise regression

***Supplementary Data 7.*** *Results of the contrast discrimination task*

Trials for which subjects failed to provide a response within 3000 ms were discarded from the analyses (0.5% of trials performed by subjects in the intention discrimination group and 0.3% of trials performed by subjects in the kinematic discrimination group). Furthermore, the first 25% of trials of each block were discarded, as for all analyses performed for the intention discrimination task and the kinematic discrimination task. Task performance revealed no influence of IPL cTBS or IFG cTBS in either task (*p* > 0.05).

### Supplementary Figures

**Supplementary Fig. 1.**
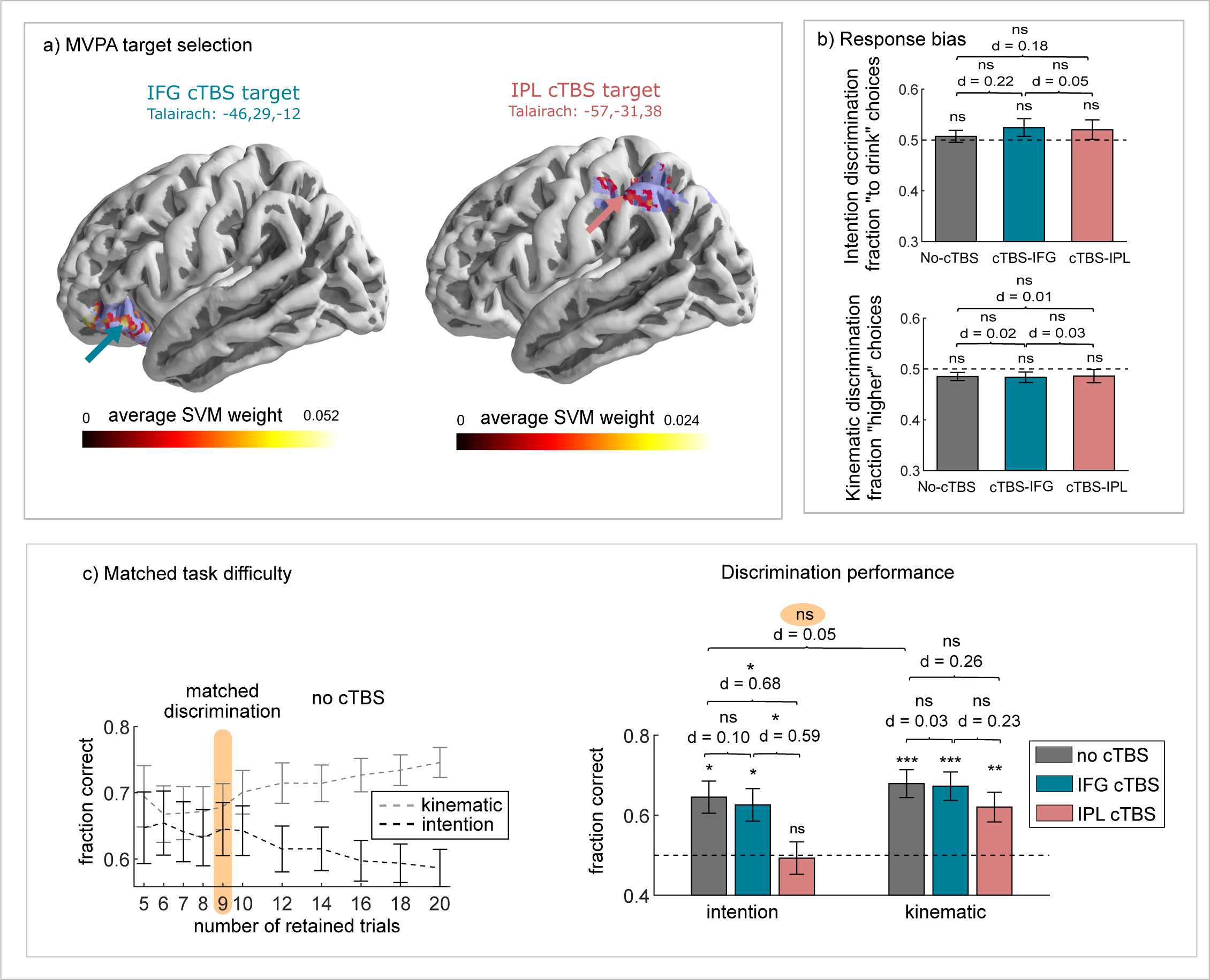
Experimental design and further analyses of behavioral discrimination performance. Related to Fig. 1. **a** Target sites of cTBS in the left IFG pars orbitalis and the left IPL anterior. Regions in slate blue indicate ROIs included in linear SVM classifier fitting. Cortical surface projections of the 20% highest ranked voxels for classifying intention in the left IFG pars orbitalis and left IPL are displayed. The displayed weights of voxels were first averaged over LOSO folds, then smoothed using a 2mm full width at half maximum (FWHM) Gaussian kernel. Arrows indicate the coordinates chosen for cTBS. **b** Fraction of ‘to drink’ answers in the intention discrimination task and ‘higher’ answers the kinematic discrimination task in each experimental session. Results are reported as mean ± SEM across subjects. ***c*** Control analyses with matched discrimination performance. (Left) Discrimination performance as a function of the number of retained trials. Orange area indicates the 10% level trial selection. With this selection, discrimination performance did not differ between the two tasks in no cTBS. (Right) Discrimination performance (fraction correct) in the intention discrimination task and in the kinematic discrimination task with 10% level trial selection. Histograms represent mean ± SEM across participants. Cohen’s effect size (d) for each comparison is reported.

**Supplementary Fig. 2.**
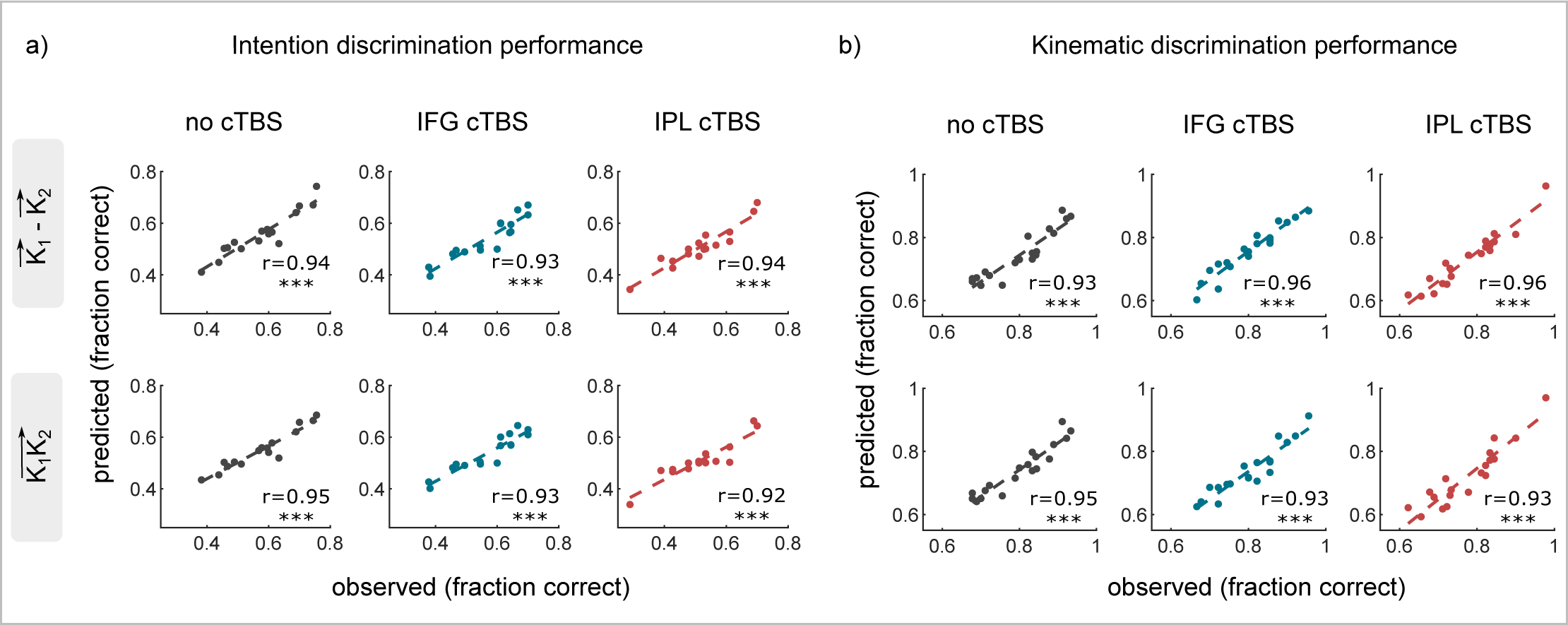
Comparison between models using the difference in kinematic features and models using all the kinematic features of the two reach-to-grasp acts. Related to Fig. 3. **a-b** Scatterplot of the relationship between the observed discrimination performance and the one predicted by the readout model across individual participants in the intention discrimination task (a) and in the kinematic discrimination task (b).

**Supplementary Fig. 3.**
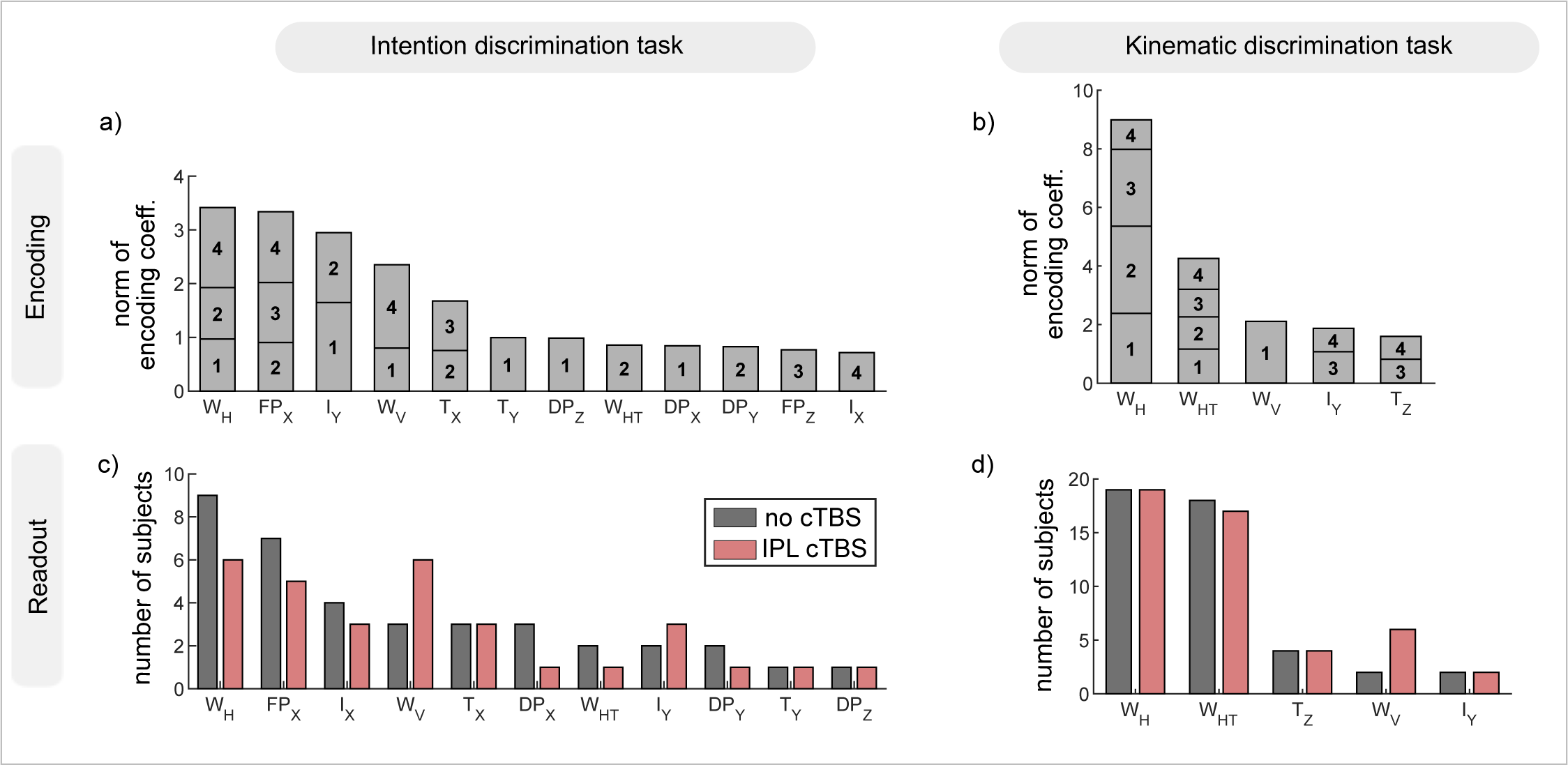
Ranking of features with respect to encoding and readout. Related to Fig. 5. **a-b** Sum of the absolute value of encoding regression coefficients over time bins encoding significant intention (a) and wrist height (b) discriminative information. Features are ordered left to right from most informative to least informative. **c-d** Number of observers who read out a specific variable in any of the time bins in the intention discrimination task (c) and in the kinematic discrimination task (d). Features are ordered left to right from most to least readout.

**Supplementary Fig. 4.**
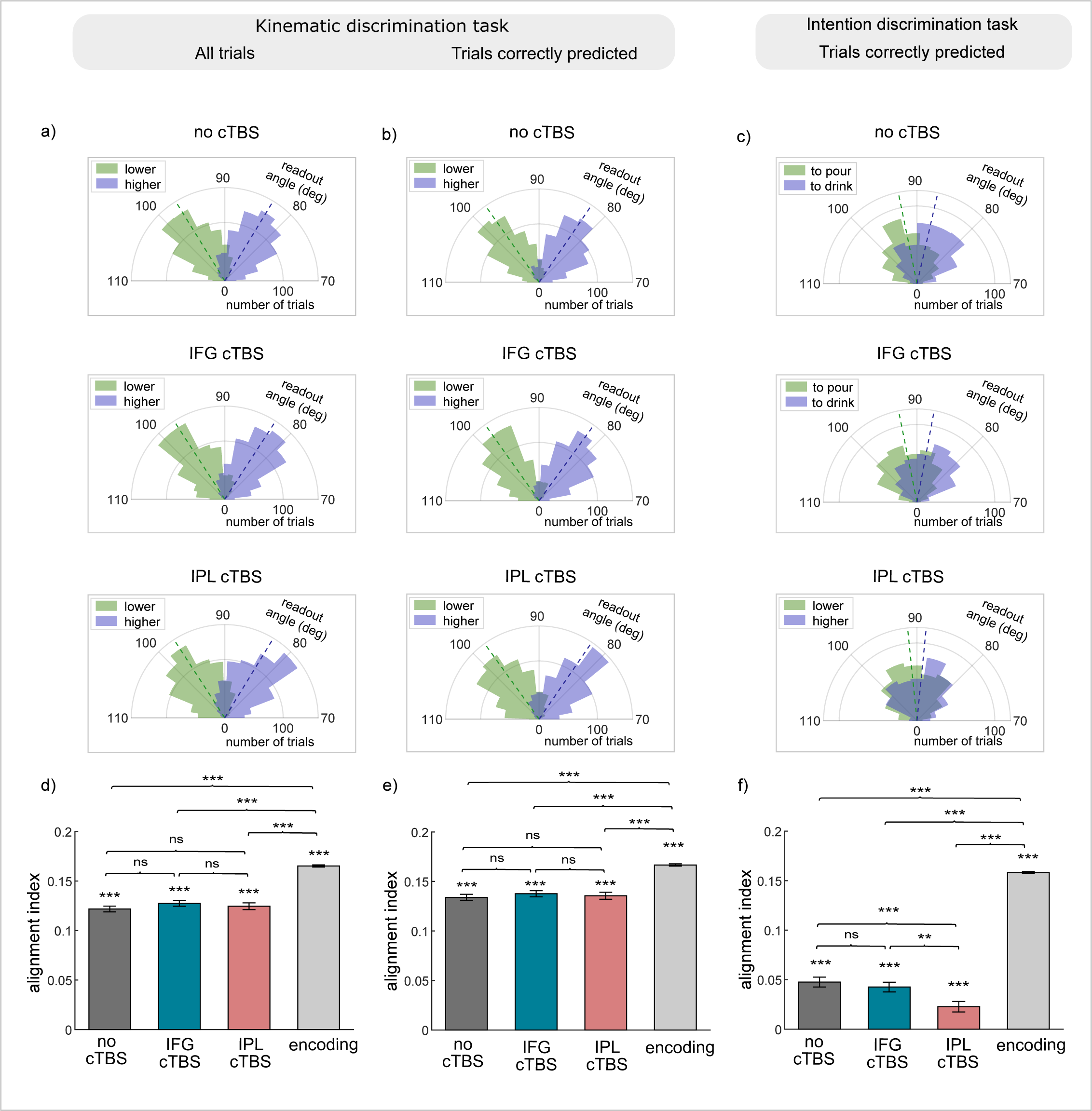
Additional analyses on the effect of cTBS on alignment. Related to Fig. 4. **a-b** Polar distribution of readout angles for ‘higher’ and ‘lower’ trials under no cTBS, IFG cTBS and IPL cTBS conditions in the kinematic discrimination task. Panel (a) reports results for all trials. Panel (b) reports results for trials correctly predicted by the model only. **c** Polar distribution of readout angles for ‘to pour’ and ‘to drink’ trials under no cTBS, IFG cTBS and IPL cTBS in the intention discrimination task considering trials correctly predicted by the model only. For graphical representation, in panels (a)-(c), the 70-110° angle range of polar distributions are expanded to a semi-circle. **d-e**. Effect of cTBS on the alignment index the kinematic discrimination task. Panel (d) reports results for all trials. Panel (e) reports results for trials correctly predicted by the model only. **f** Effect of cTBS on the alignment index the intention discrimination task considering trials correctly predicted by the model only. In panels (d)-(f), the value of the alignment index of the encoding angle is also reported for comparison. Histograms represent mean ± SEM across all trials and participants.

**Supplementary Fig. 5.**
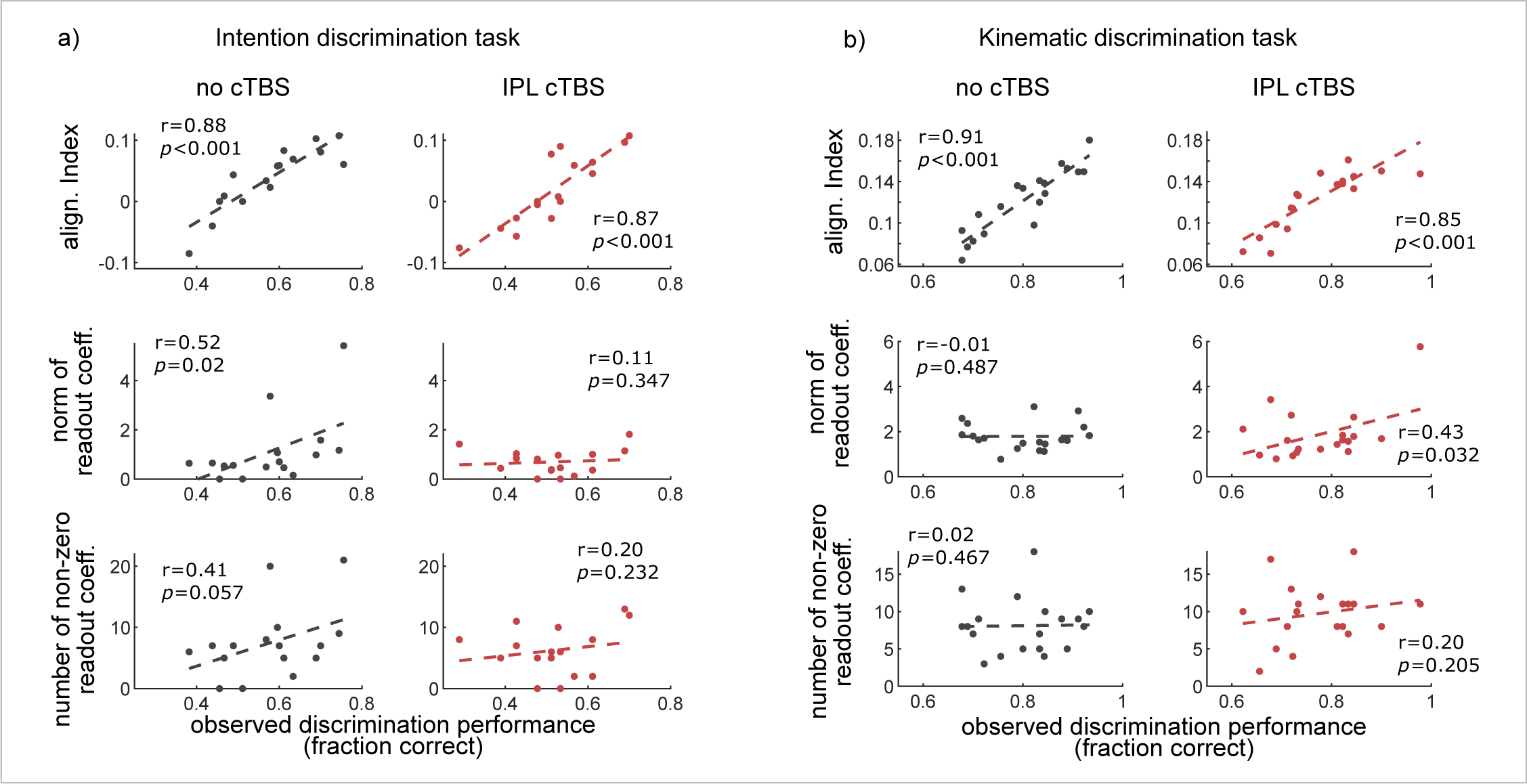
Correlation of model parameters with task performance. Related to Fig. 4. ***a-b*** Scatterplot of the alignment index, the norm of readout vector and the number of non-zero readout coefficients against observed discrimination performance across participants under no cTBS and IPL cTBS sessions in the intention discrimination task (a) and in the kinematic discrimination task (b).

**Supplementary Fig. 6.**
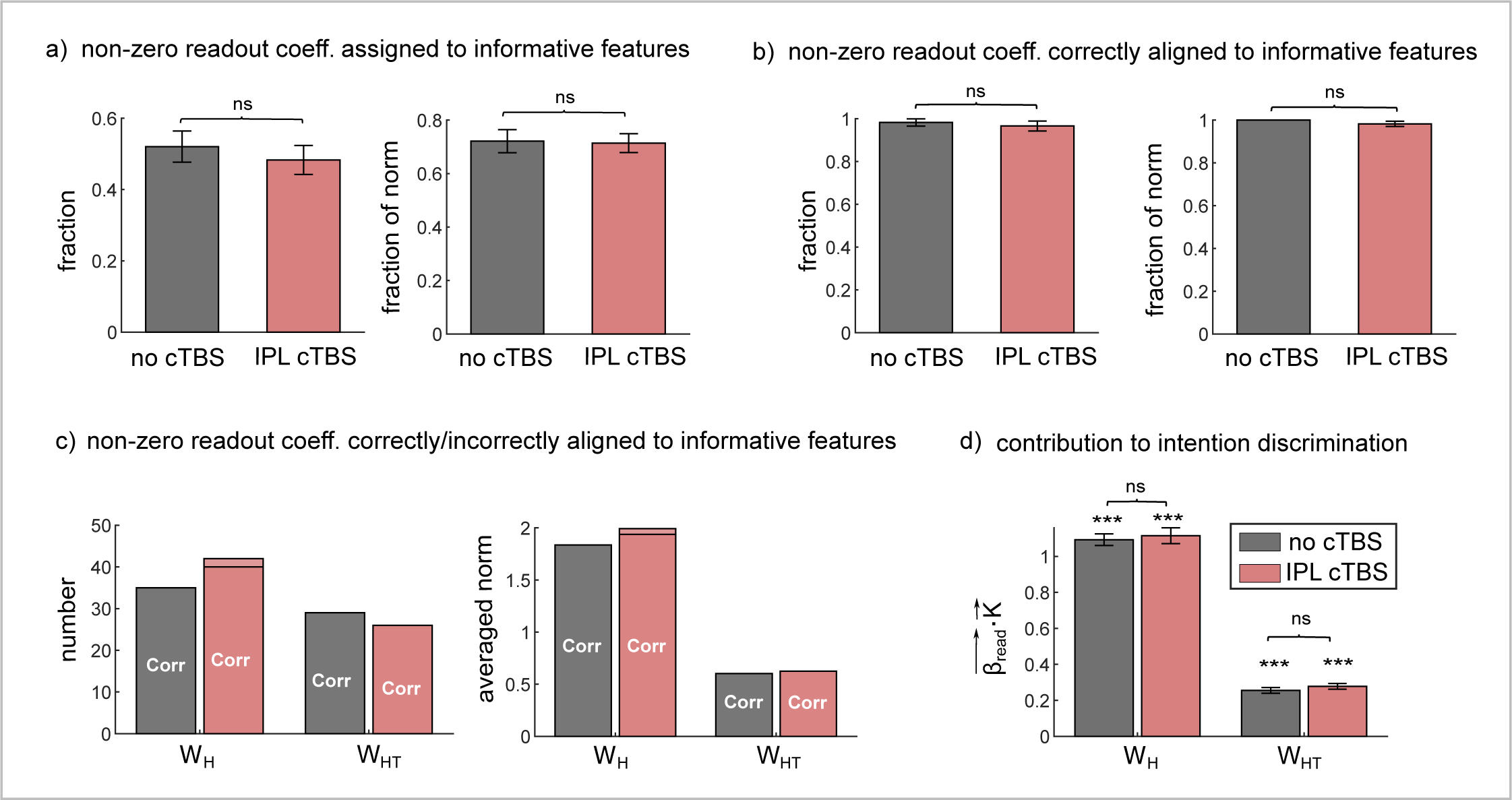
Quantification of alignment in the kinematic discrimination task. **a** Fraction and fraction of the norm of non-zero readout coefficients assigned to informative features. **b** Fraction and fraction of the norm of non-zero readout coefficients assigned to informative features and correctly aligned with encoding. Fraction was computed on a subject basis and then averaged across subjects. **c** Number and averaged norm of non-zero readout coefficients (in)correctly aligned to informative features in encoding. We focused on the most informative and most read out kinematic variable: the height of the wrist (W_H_) and the horizontal trajectory of the wrist (W_HT_). **d** Contribution of W_H_ and W_HT_ to kinematic discrimination performance, computed as the scalar product between the kinematic vector and the readout vector within that feature subspace. Histograms represent mean ± SEM across all trials and participants.

